# A general exposome factor explains individual differences in functional brain network topography and cognition in youth

**DOI:** 10.1101/2023.08.25.554893

**Authors:** Arielle S. Keller, Tyler M. Moore, Audrey Luo, Elina Visoki, Mārtiņš M. Gataviņš, Alisha Shetty, Zaixu Cui, Yong Fan, Eric Feczko, Audrey Houghton, Hongming Li, Allyson P. Mackey, Oscar Miranda-Dominguez, Adam Pines, Russell T. Shinohara, Kevin Y. Sun, Damien A. Fair, Theodore D. Satterthwaite, Ran Barzilay

## Abstract

Our minds and brains are highly unique. Despite the long-recognized importance of the environment in shaping individual differences in cognitive neurodevelopment, only with the combination of deep phenotyping approaches and the availability of large-scale datasets have we been able to more comprehensively characterize the many inter-connected features of an individual’s environment and experience (“exposome”). Moreover, despite clear evidence that brain organization is highly individualized, most neuroimaging studies still rely on group atlases to define functional networks, smearing away inter-individual variation in the spatial layout of functional networks across the cortex (“functional topography”). Here, we leverage the largest longitudinal study of brain and behavior development in the United States to investigate how an individual’s exposome may contribute to functional brain network organization leading to differences in cognitive functioning. To do so, we apply three previously-validated data driven computational models to characterize an individual’s multidimensional exposome, define individual-specific maps of functional brain networks, and measure cognitive functioning across broad domains. In pre-registered analyses replicated across matched discovery (*n*=5,139, 48.5% female) and replication (*n*=5,137, 47.1% female) samples, we find that a child’s exposome is associated with multiple domains of cognitive functioning both at baseline assessment and two years later – over and above associations with baseline cognition. Cross-validated ridge regression models reveal that the exposome is reflected in children’s unique patterns of functional topography. Finally, we uncover both shared and unique contributions of the exposome and functional topography to cognitive abilities, finding that models trained on a single variable capturing a child’s exposome can more accurately and parsimoniously predict future cognitive performance than models trained on a wealth of personalized neuroimaging data. This study advances our understanding of how childhood environments contribute to unique patterns of functional brain organization and variability in cognitive abilities.

## INTRODUCTION

Our minds and brains are highly unique, shaped not just by our genetics (“genome”) but also by our complex network of environmental exposures (“exposome”). In line with this idea is the observation that individual differences in cognition tend to increase throughout development (Kidd et al., 2018) as environmental exposures and individual experiences continually mold the brain’s functional organization. It is imperative that we characterize how individual differences in cognition emerge during childhood, not only as a window into understanding what makes us who we are as unique individuals, but also because individual differences in cognitive functioning are associated with critical socio-economic (Cortés Pascual et al., 2019; Moffitt et al., 2011), physical health (Batty et al., 2007; Calvin et al., 2011; Gow et al., 2011; Hart et al., 2004; Wraw et al., 2015), and mental health (Barzilay et al., 2019; Klassen et al., 2004; Shamosh et al., 2008; Shanmugan et al., 2016) outcomes across the lifespan. Understanding cognitive neurodevelopment at the level of the individual therefore requires a characterization of how the unique features of each child’s environment may be reflected in the unique features of each child’s individual brain.

Many aspects of a child’s environment and experiences have been linked with cognitive functioning in previous work. The Adolescent Brain Cognitive Development (ABCD) Study (Volkow et al., 2018) (*n*=11,878) is an ongoing large-scale longitudinal study of development with deep phenotyping of multidimensional environmental features and cognitive abilities in children from twenty-two sites across the United States. In the ABCD Study, environmental features that have been shown to be associated with cognitive functioning include family dynamics (Gong et al., 2021; Thompson et al., 2022; Zhang et al., 2020), experiences at school (Meredith et al., 2022; Rakesh et al., 2021) and online (Kirlic et al., 2021; Paulus et al., 2019; Sauce et al., 2022), socio-economic status (Botdorf et al., 2022; Dennis et al., 2022), physical activity (Ronan et al., 2020; Walsh et al., 2018), prenatal substance use (Cioffredi et al., 2022) and stress (Demidenko et al., 2021). The diverse features of a child’s environment that may independently and interactively influence cognitive neurodevelopment pose a challenge for studies aiming to characterize these important relationships by necessitating both large sample sizes and deep phenotyping. Despite the long-recognized importance of the environment in shaping cognitive development, only recently have we been able to leverage data-driven approaches in deeply phenotyped datasets of this size to capture the multitude of inter-connected features that make up an individual’s environment and experience.

The concept of the “exposome” has been introduced as a way of capturing the totality of environmental exposures and experiences throughout an individual’s lifespan (Rappaport, 2011; Wild, 2005). While the first exposome studies primarily focused on associations with physical health (e.g., cancer risk) in adults (Rappaport, 2011; Wild, 2005), more recent work has increased focus on mental health outcomes (Guloksuz et al., 2018) including recent studies turning this focus to children. Factor analytic approaches have made it possible to quantify both a child’s general exposome as well as identify sub-factors capturing specific aspects of a child’s perinatal, familial, social, and physical environments (Moore et al., 2022), all of which can have substantial effects on mental functioning. This approach has revealed that a child’s exposome is associated with psychopathology (Barzilay et al., 2022; Moore et al., 2022; Pries et al., 2022).

In a similar vein, an increase in large-scale deeply phenotyped neuroimaging datasets have made it possible to define robust multidimensional features of an individual’s functional brain organization at scale. Numerous studies have highlighted the striking inter-individual heterogeneity in the size, shape and spatial arrangement of functional brain regions across the cortex (Laumann et al., 2015; Gordon et al., 2017; Glasser et al., 2016; Kong et al., 2019; Li et al., 2017; Bijsterbosch et al., 2019). Despite this well-documented heterogeneity, the majority of human neuroimaging studies still rely on standardized network atlases of functional networks (Power et al., 2011; Yeo et al., 2011) that are spatially warped to individual brains, smearing away the rich complexity of individual differences. Cognitive functions in particular are supported by spatially-distributed, large-scale association networks (Dosenbach et al., 2007; Owen et al., 2005; Corbetta & Shulman, 2002) that tend to have the *highest* inter-individual heterogeneity among the many functional networks of the human connectome in both adults (Gordon et al., 2017; Gratton et al. 2018) and kids (Cui et al., 2020), exacerbating this problem for studies of cognitive development. To overcome this challenge, precision functional mapping techniques have been developed as an alternative to group-level atlases, deriving individually-defined networks that capture each brain’s unique pattern of functional topography. Such personalized functional networks (PFNs) have been found to be highly stable within individuals and to predict an individual’s spatial pattern of activation on fMRI tasks (Gordon et al., 2017; Glasser et al., 2016; Tavor et al., 2016).

Here, we investigate the role of children’s complex, interconnected environments and experiences in shaping unique patterns of functional brain network topography and cognitive development. To characterize reproducible cross-sectional and longitudinal environment-brain-behavior associations, we leverage ABCD Study^Ⓡ^ data to conduct our pre-registered analyses (Keller and Barzilay, 2023). We use three previously validated data-driven approaches to reduce dimensionality across rich multivariate data types: 1) we define both general and specific exposome factors using longitudinal exploratory bifactor analysis (Moore et al., 2022); 2) we define personalized functional networks (PFNs) using non-negative matrix factorization (Cui et al., 2020; Keller et al., 2022b; Lee and Seung, 1999; Li et al., 2017); and 3) we define cognitive factors using Bayesian principal components analysis from a previous study in this dataset (Thompson et al., 2019). Using linear mixed effects models and cross-validated ridge regressions, we relate individual differences in the exposome to PFN topography and individual differences in cognitive functioning across domains. Given the critical importance of using large samples to identify reproducible brain-behavior associations (Marek, Tervo-Clemmens et al., 2022), we replicate our analyses across matched discovery (*n*=5,139, 48.5% female) and replication (*n*=5,137, 47.1% female) samples (Feczko et al., 2021; Cordova et al., 2021). Our results show that a child’s exposome is associated with multiple domains of cognitive functioning and predicts cognitive abilities two years later over and above the effect of baseline cognition. Additionally, we find that the exposome is reflected in children’s unique patterns of functional network topography, and we uncover both shared and unique contributions of the exposome and brain organization in shaping cognitive abilities. Together, these findings provide a characterization of multivariate associations among environment, brain and cognition, highlighting the critical role that childhood environments play in shaping unique patterns of functional brain organization and diverse cognitive abilities.

## RESULTS

### A general measure of the exposome is associated with individual differences in cognition across multiple tasks

We first aimed to characterize associations between a child’s complex, multidimensional environment (“exposome”) and their cognitive abilities. To do so, we derived a measurement of each child’s exposome using multilevel (clustered) exploratory factor analysis with a bifactor rotation (Jennrich and Bentler, 2011), with the same goal as that outlined in Moore et al. (2022). Given that we aimed to investigate both cross-sectional and longitudinal associations among the exposome, functional brain network topography and cognition, we derived exposome scores using data from multiple timepoints, accounting for clustering by family, stratification by site, and model constraints ensuring measurement invariance across time. Specifically, the model was constrained to have the same configuration, loadings, and intercepts (with unconstrained factor means) across time points. If this constraint were unrealistic–i.e. if the exposome models differed across time points–this would be reflected in the model fit indices, providing an embedded check on the assumption of measurement invariance across time (age). We used a bifactor approach (Reise, 2012; Reise et al., 2010) because, based on prior work (Moore et al., 2022), we anticipated that a general exposome factor would capture variance across dimensions of a child’s complex environment (e.g., neighborhood, school, etc.; **Figure 1a**). In addition to a general exposome factor score for each participant, the bifactor model provided six sub-factors capturing specific dimensions of a child’s environment: School, Family Values, Family Turmoil, Dense Urban Poverty, Extracurriculars and Screen Time. These sub-factors are necessarily orthogonal to one another as well as orthogonal to the general exposome factor (**Supplementary** Figure 1). As depicted in **Supplementary Table 1**, the variables loading most strongly on the general exposome factor were those capturing dimensions of socio-economic status (e.g., household income, parental education and marital status, children’s involvement in extracurricular sports/activities, neighborhood safety, crowding and crime), with positive associations between general exposome scores and socio-economic status (**Supplementary** Figure 2).

**Figure 1.**
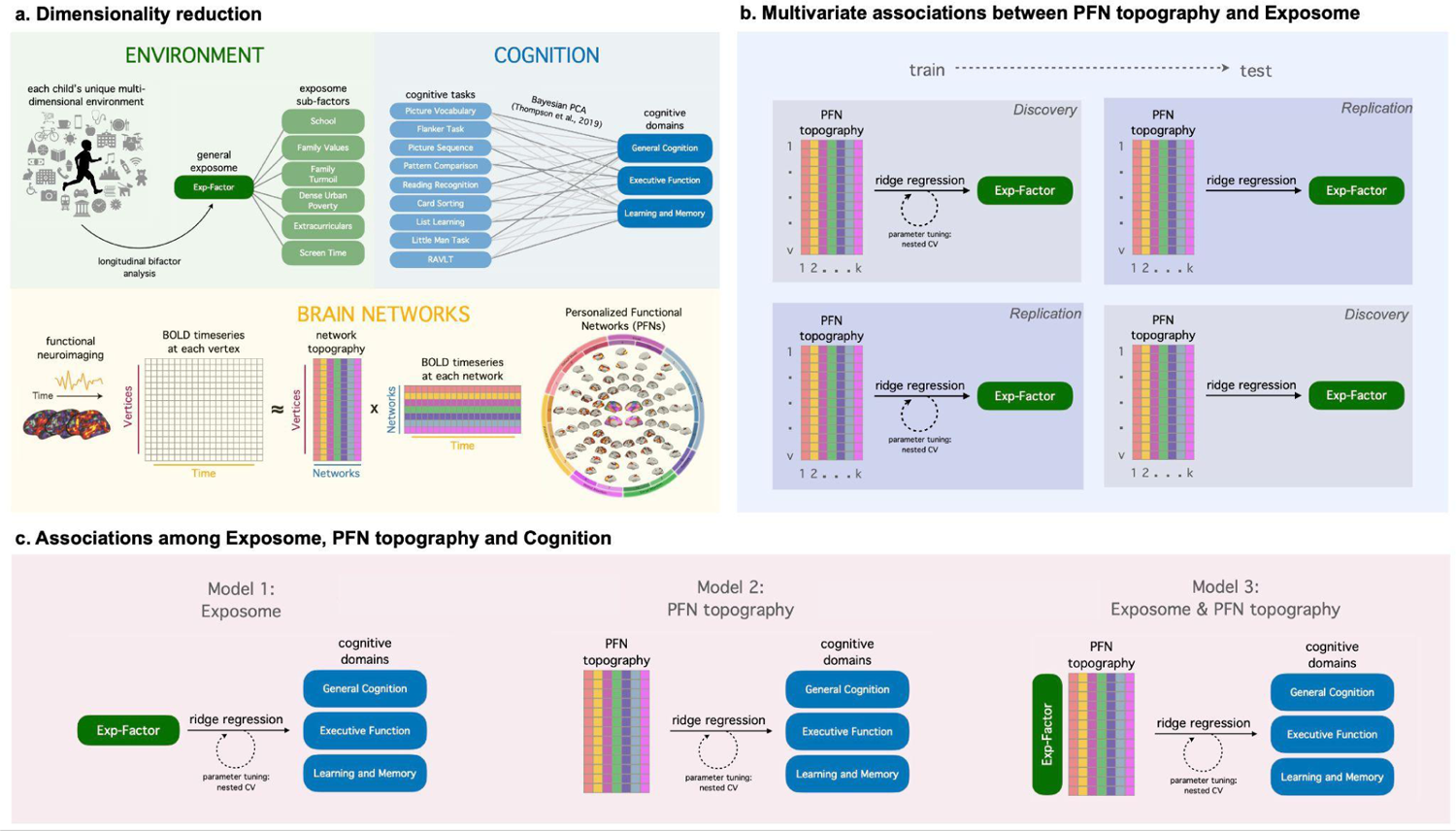
Summary of Methodological Approach. **a** We aimed to define broad factors describing a child’s environment, cognition, and brain network organization by reducing high-dimensional data with three unsupervised machine-learning methods. To identify general and specific factors of each child’s complex multidimensional environment, we began with a set of 354 variables capturing a variety of features and applied longitudinal exploratory bifactor analysis with a goal analogous to our previous work (Moore et al., 2022). To capture three broad domains of cognition across nine well-validated cognitive tasks, we used the results of a Bayesian probabilistic principal components analysis from a previous study in this dataset (Thompson et al., 2019). To map the spatial topography of personalized functional networks (PFNs) within each child’s brain, we assessed seventeen PFNs identified by spatially-regularized non-negative matrix factorization of each child’s functional connectome as derived in our prior work (Keller et al., 2022; Cui et al., 2020; Li et al., 2017). **b** To investigate whether children’s general exposome is encoded in their multivariate patterns of PFN topography, we trained ridge regression models using two-fold cross-validation across our matched discovery and replication samples. This involved first training a model in the discovery dataset and testing its out-of-sample performance in the replication dataset, and then reversing the order (training in the replication dataset and testing in the discovery dataset) in order to assess model performance on held-out data across the entire sample. All ridge regression models accounted for covariates for age, biological sex, scanning site, and head motion, and the ridge parameter was tuned using nested cross-validation during the training phase to avoid overfitting. To ensure that model performance was not influenced by the choice of split, we also performed repeated random cross-validation using multiple random splits and found comparable results. **c** To investigate associations among the exposome, PFN topography, and cognition, we trained three models using the same procedure as above: Model 1 (“Exposome”) used only a participant’s general exposome score; Model 2 (“PFN Topography”), reported in our prior work (Keller et al., 2022b), used each participant’s multivariate pattern of personalized functional network topography; and Model 3 (“Exposome + PFN Topography”) used a combination of the features in the first and second model types, hypothesized to perform best by capitalizing on both shared and unique variance across features. By evaluating model performance and comparing performance among the three model types, we were able to determine the relative contributions of a child’s exposome and functional brain network organization to cognitive functioning.

To investigate associations between exposome factor scores and cognition at baseline (9-10 years old) and two-year follow-up (11-12 years old), we estimated cross-sectional and longitudinal linear mixed effects models. All models accounted for family structure and ABCD Study site as random effects as well as age and biological sex as fixed effects. Given that the general exposome factor and six exposome sub-factors are necessarily orthogonal (**Supplementary** Figure 1), we included all seven factors together in our models without violating assumptions of collinearity. Across demographically-matched discovery (*n*=5,139, 48.5% female) and replication (*n*=5,137, 47.1% female) samples (**Supplementary Table 2**), the general exposome factor was significantly associated with cognition on all five cognitive tasks at baseline (Figure 2). These associations remained significant with the inclusion of all six exposome sub-factor scores in addition to age, sex, site, and family covariates and survived Bonferroni correction for multiple comparisons (discovery: *r*s=0.12-0.50, all *p_bonf_*<.001; replication: *r*s=0.15-0.48, all *p_bonf_*<.001; **Supplementary Table 3**). Notably, these associations also remained significant in longitudinal models predicting cognition two years later while accounting for baseline cognition, which is known to be a strong predictor of future cognitive performance (discovery: *r*s=0.08-0.24, all *p_bonf_*<.001; replication: *r*s=0.11-0.22, all *p_bonf_*<.001; **Table 1**). These longitudinal results demonstrate that the general exposome factor measured at ages 9-10 could predict cognitive performance two years later when children were 11-12 years old. Sensitivity analyses revealed that these results were also significant over and above the effects of commonly utilized measures of socio-economic status (household income and parental education) as well as over and above the effects of psychiatric medication use (ADHD medications, antipsychotics and antidepressants) (**Supplementary Tables 4 and 5**). Stratified analyses by racial identification and biological sex revealed consistent positive associations between the general exposome factor and cognitive functioning for nearly all cognitive tasks in all groups, with only a small subset of these associations not passing correction for multiple comparisons (**Supplementary Table 6**).

**Figure 2.**
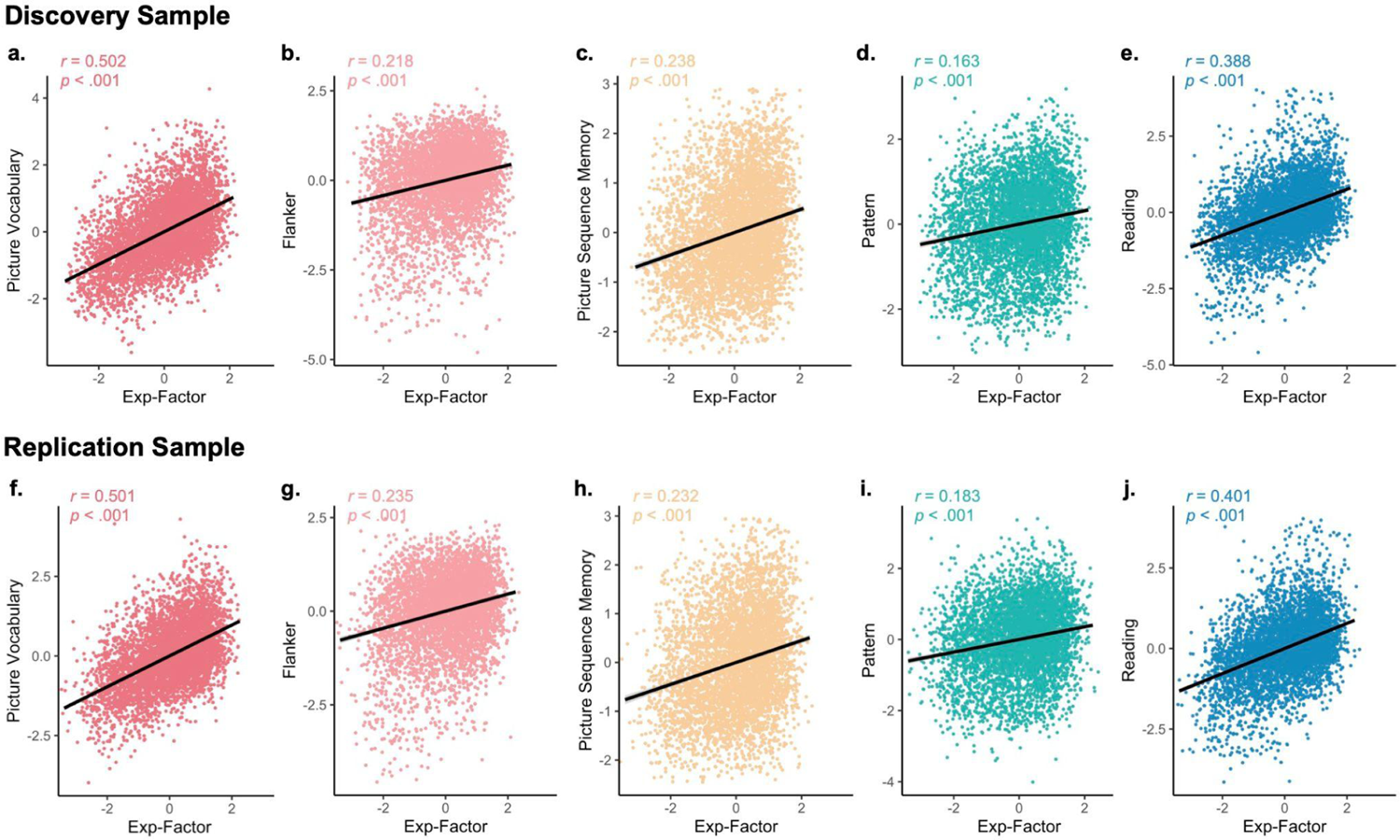
A general measure of exposome is significantly associated with cognition across cognitive tasks. We characterized a child’s unique, complex and multidimensional exposome using a general exposome factor (“Exp-Factor”) derived from longitudinal bifactor analysis. The general exposome factor is significantly associated with cognitive performance across all five tasks assessed across both the discovery (**a-e**) and replication (**f-j**) samples. One outlier was excluded from Panel A for visualization purposes (Picture Vocabulary score less than −4) but this participant was retained for all statistical analyses.

**Table 1.**
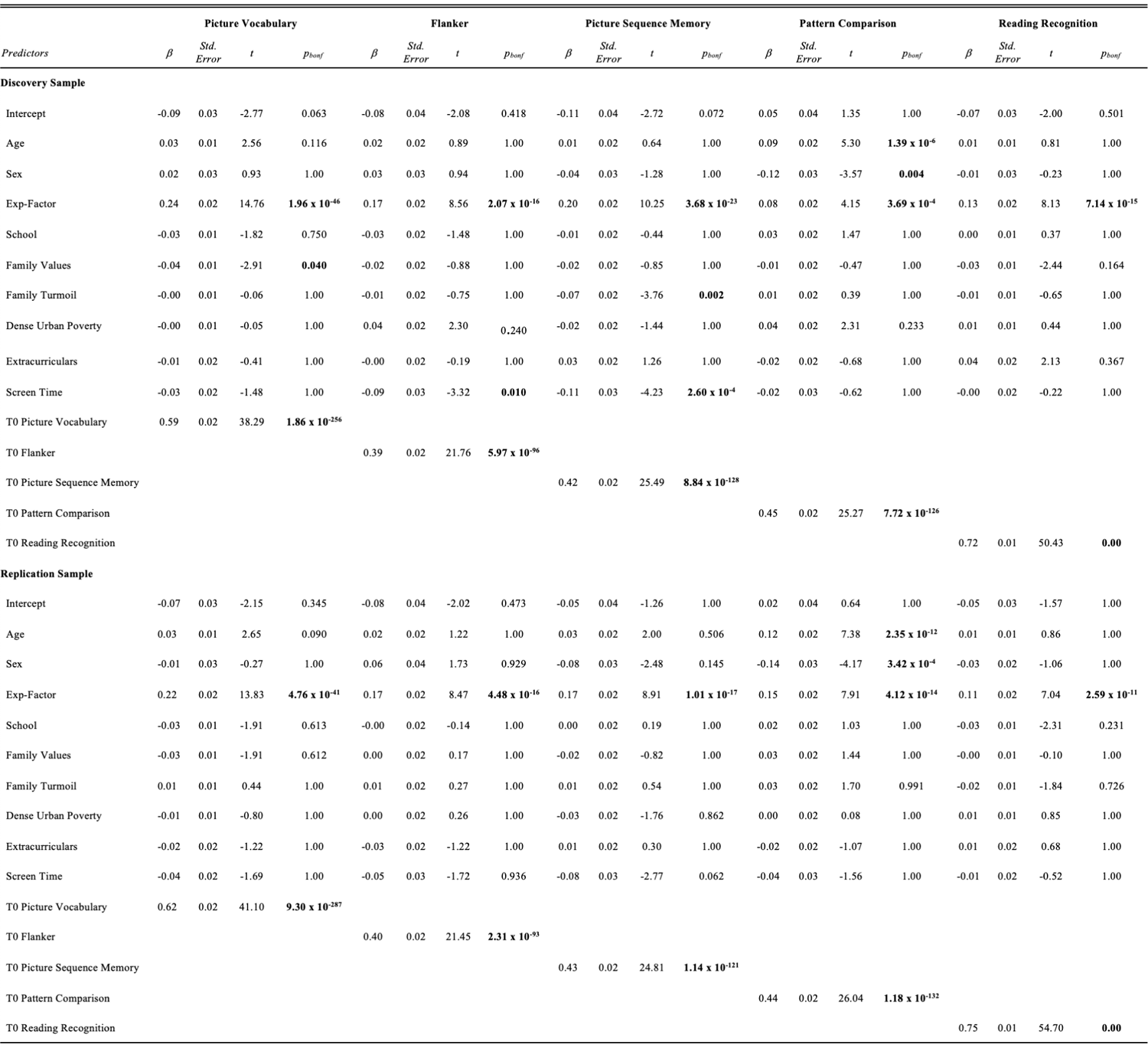
The general exposome factor at ages 9-10 is significantly associated with cognition at ages 11-12 over and above baseline cognitive performance. Across all five cognitive tasks and across both the discovery and replication sub-samples, the general exposome factor (“Exp-Factor”) is positively associated with cognition. These effects held with the inclusion of baseline (T0) cognitive performance. Note that random effects for site and family are also included as covariates in these models.

### Exposome scores are reflected in the multivariate pattern of personalized functional brain network topography in youth

Having demonstrated that exposome scores are strongly associated with cognitive functioning in youth, we next sought to understand the extent to which a child’s general exposome contributes to their individual functional brain network organization. To capture the profound inter-individual heterogeneity in the spatial topography of functional brain networks in children, we defined a unique map of functional brain networks for each child using non-negative matrix factorization as previously reported (Figure 1a; Cui et al., 2020; Keller et al., 2022b). Personalized functional network (PFN) topography was defined as each individual’s multivariate pattern of vertex-wise loadings for each of 17 PFNs. To investigate associations between these high-dimensional multivariate patterns of PFN topography and each child’s general exposome factor, we trained ridge regression models using two-fold cross-validation across our matched discovery and replication samples (Figure 1b) and only report results from testing our models on unseen data. Here, the ridge parameter (L2-norm) was tuned by nested cross-validation to avoid overfitting and then applied to weight PFN loadings by the strength of their associations with the outcome variable, reducing model complexity. All ridge regression models accounted for covariates for age, biological sex, scanning site, and head motion (mean framewise displacement).

PFN topography was associated with the general exposome factor in unseen data, with significant correlations between a child’s observed general exposome factor score and the general exposome factor score estimated by the ridge regression models (Figure 3a, discovery: *r* = 0.440, *p*<0.001, 95% CI: [0.41, 0.47]; replication: *r* = 0.462, *p*<0.001, 95% CI: [0.44, 0.49]).

**Figure 3.**
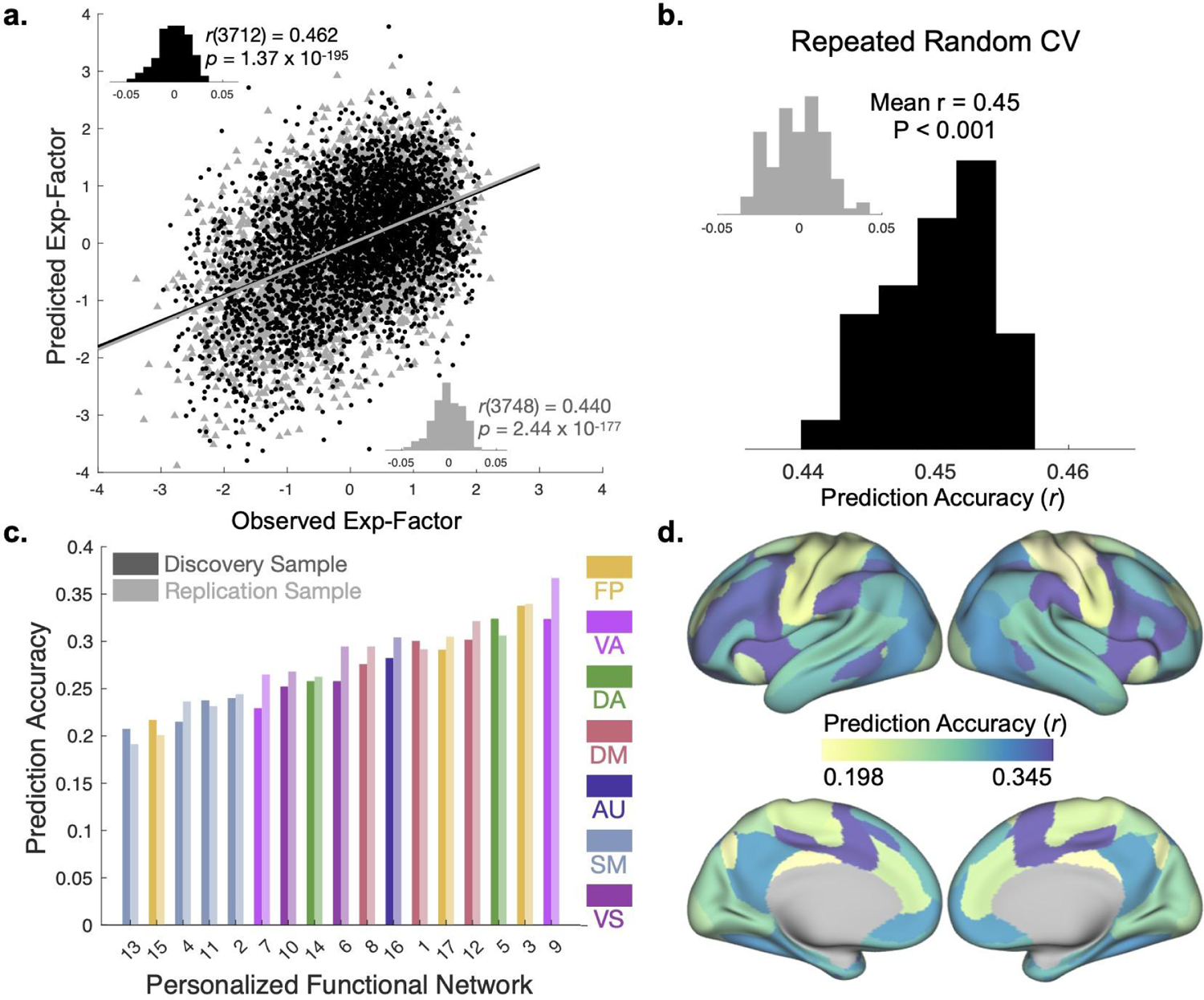
Exposome scores are reflected in the multivariate pattern of personalized functional brain network topography. Prediction accuracy is defined as the correlation between observed general exposome factor (calculated based on the child’s individual environment from various measurements in the ABCD Study^Ⓡ^) and the general exposome factor score that was derived from ridge regression models trained on PFN functional topography**. a** Association between observed and predicted general exposome factor scores using two-fold cross-validation (2F-CV) with nested cross-validation for parameter tuning across both the discovery (black scatterplot) and replication (gray scatterplot) samples. Inset histograms represent null distributions of prediction accuracies with permuted data. **b** Repeated random 2F-CV (100 runs) provided evidence of stable prediction accuracy across splits of the data, which was far better than a null distribution with permuted data (inset). **c** Independent network models reveal that association networks such as the fronto-parietal and ventral attention networks yielded the most accurate predictions of the general exposome factor. Note that all *p*-values associated with prediction accuracies are significant after Bonferonni correction for multiple comparisons. Numerical assignments for each PFN are consistent with previous work (Cui et al., 2020; Keller et al., 2022). **d** Prediction accuracy averaged across discovery and replication samples depicted for seventeen cross-validated models trained on each PFN independently. *Abbreviations:* Exp-Factor: general exposome factor; CV: cross-validation; FP: fronto-parietal; VA: ventral attention; DA: dorsal attention; DM: default mode; AU: auditory; SM: somatomotor; VS: visual.

Repeated random cross-validation confirmed that our results were not driven by the choice of split across matched discovery and replication samples (Figure 3b; mean *r* = 0.45, p<.001). To further confirm that our results were not driven by leakage across samples, we repeated this same model training and testing procedure using exposome scores that were generated independently in the discovery and replication samples rather than from the full sample of participants, and found nearly identical results (**Supplementary** Figure 3, discovery: *r* = 0.440, *p*<0.001, 95% CI: [0.41, 0.47]; replication: *r* = 0.460, *p*<0.001, 95% CI: [0.43, 0.49]). Independent ridge regression models trained on the functional topography of each of the 17 PFNs separately highlight the distribution of prediction accuracies (correlations between true general exposome factor and model-generated general exposome factor) across networks: association networks (e.g., fronto-parietal networks) and attention networks (e.g., dorsal and ventral attention networks) yielded higher accuracy while sensorimotor networks (e.g., somatomotor networks) yielded lower accuracy (Figure 3c**,d**). Together, these findings reveal a clear association between a child’s exposome score and their unique multivariate pattern of PFN topography.

### Exposome scores and personalized functional brain network topography are associated with cognition

To compare multivariate associations among exposome scores, personalized functional brain network topography and cognitive functioning, we trained three types of linear ridge regression models to predict three domains of cognition (General Cognition, Executive Function, and Learning/Memory). The first model type (“Exp-Factor”) used only a participant’s general exposome score, while the second model type (“PFN Topography”), reported in our prior work (Keller et al., 2022b), used each participant’s multivariate pattern of personalized functional network topography. The third model type (“Exp-Factor + PFN Topography”) used a combination of the features in the first and second model types, hypothesized to perform best by capitalizing on both shared and unique variance across features. By evaluating model performance and comparing performance among the three model types, we were able to determine the relative contributions of a child’s experiences/environment and their complex, individualized pattern of functional brain network organization to cognitive functioning. Note that the model training procedure with ridge regression remained the same even for the single-variable model to allow the slope of the association between the general exposome factor and cognition to vary with the value of the ridge constraint during nested cross-validation for parameter tuning. All three models performed well, yielding highly significant correlations between a child’s true cognitive performance scores and the model-generated cognitive performance scores (**Table 2**), with the highest accuracy for predictions of General Cognition and lower accuracy for the Learning/Memory and Executive Function domains.

**Table 2.** Comparison of models relating exposome and PFN topography to cognitive functioning across domains. To compare multivariate associations among exposome scores, personalized functional brain network topography and cognitive functioning, we trained linear ridge regression models to predict three domains of cognition (General Cognition, Executive Function, and Learning/Memory). The first model type (“Exp-Factor”) used only a participant’s general exposome score, while the second model type (“PFN Topography”), reported in our prior work (Keller et al., 2022b), used each participant’s multivariate pattern of personalized functional network topography. The third model type (“Exp-Factor + PFN Topography”) used a combination of the features in the first and second model types. Correlations between true cognitive performance and model-generated cognitive performance (*r*) were significant for all model types and highest for the predictions of General Cognition. Model comparison using the Akaike Information Criterion (AIC) and Bayesian Information Criterion (BIC), indices that account for differences in the number of features across models, reveals that the Exp-Factor model is the most parsimonious.

| *Prediction Accuracy* | **Discovery** |  |  |  | **Replication** |  |  |  |
| --- | --- | --- | --- | --- | --- | --- | --- | --- |
|  | *r* | *p* | AIC | BIC | *r* | *p* | AIC | BIC |
| **General Cognition** |  |  |  |  |  |  |  |  |
| Exp-Factor | 0.42 | $2.65 \times 10^{-148}$ | -248.8198 | -8.0250 | 0.46 | $1.17 \times 10^{-179}$ | -454.2790 | -7.8809 |
| PFN Topography | 0.41 | $3.05 \times 10^{-146}$ | $2.0198 \times 10^6$ | $8.2493 \times 10^6$ | 0.45 | $3.85 \times 10^{-174}$ | $2.0196 \times 10^6$ | $8.2267 \times 10^6$ |
| Exp-Factor + PFN Topography | 0.44 | $2.44 \times 10^{-166}$ | $2.0196 \times 10^6$ | $8.2491 \times 10^6$ | 0.48 | $3.30 \times 10^{-194}$ | $2.0195 \times 10^6$ | $8.2270 \times 10^6$ |
| **Executive Function** |  |  |  |  |  |  |  |  |
| Exp-Factor | 0.11 | $8.86 \times 10^{-11}$ | 1217.1946 | -8.8570 | 0.14 | $7.30 \times 10^{-16}$ | 1035.4933 | -8.7450 |
| PFN Topography | 0.17 | $1.37 \times 10^{-23}$ | $2.0210 \times 10^6$ | $8.2493 \times 10^6$ | 0.16 | $5.48 \times 10^{-22}$ | $2.0209 \times 10^6$ | $8.2267 \times 10^6$ |
| Exp-Factor + PFN Topography | 0.17 | $4.41 \times 10^{-24}$ | $2.0210 \times 10^6$ | $8.2491 \times 10^6$ | 0.17 | $8.54 \times 10^{-23}$ | $2.0209 \times 10^6$ | $8.2270 \times 10^6$ |
| **Learning/Memory** |  |  |  |  |  |  |  |  |
| Exp-Factor | 0.25 | $1.35 \times 10^{-50}$ | 470.3910 | -8.4332 | 0.27 | $6.20 \times 10^{-57}$ | 386.2874 | -8.3685 |
| PFN Topography | 0.27 | $2.06 \times 10^{-61}$ | $2.0204 \times 10^6$ | $8.2493 \times 10^6$ | 0.27 | $2.91 \times 10^{-57}$ | $2.0204 \times 10^6$ | $8.2267 \times 10^6$ |
| Exp-Factor + PFN Topography | 0.28 | $3.49 \times 10^{-66}$ | $2.0203 \times 10^6$ | $8.2491 \times 10^6$ | 0.28 | $4.92 \times 10^{-63}$ | $2.0203 \times 10^6$ | $8.2270 \times 10^6$ |

We then compared performance across models using the Akaike Information Criterion (AIC) and Bayesian Information Criterion (BIC). These indices were selected to account for the substantial differences in the number of features used to train each type of model, balancing the tradeoff between model accuracy and model complexity. These model comparisons reveal that the general exposome factor model is the most parsimonious (lowest AIC and BIC), reflecting its high accuracy and low complexity. While the full Exposome + PFN Topography model yields a small increase in accuracy, this benefit is outweighed by the substantial increase in model complexity from a single feature to thousands of features. These results suggest that a single environmental variable capturing a child’s multidimensional exposome could tell us just as much (or more) about a child’s cognitive abilities as do a wealth of robust, personalized neuroimaging variables capturing each child’s personalized functional brain network organization.

To determine which model type could best predict *future* cognitive performance, we trained ridge regression models on baseline (9-10 years old) general exposome factor and PFN topography to predict cognition assessed two years later (11-12 years old). All models were trained in the same manner as described above, with the exception that baseline cognitive performance scores for each task were included as covariates. Again, we found that the general exposome factor models were the most parsimonious across all five cognitive tasks (**Supplementary Table 7**). This finding further highlights the importance of a child’s multidimensional environment, not just for understanding current cognitive abilities but also for predicting future cognitive abilities.

Furthermore, the modest boost in model performance for the full Exp-Factor + PFN topography model compared with the PFN topography and Exp-Factor models indicates that there is a substantial amount of shared variance between the general exposome factor and PFN topography (in line with our finding that the general exposome factor is reflected in PFN topography), but there is also some unique variance explained from each feature type (see **Supplementary** Figure 4 for a comparison of prediction accuracy maps across models). To further disentangle the shared and unique variance among the exposome, functional brain network organization and cognition, we trained an additional model associating the multivariate pattern of PFN topography with general cognition after regressing out the shared variance between general cognition and the general exposome factor. This model achieved moderate accuracy (discovery: *r* = 0.240, *p*<0.001, replication: *r* = 0.233, *p*<0.001), suggesting that some features of the multivariate pattern of PFN topography are uniquely associated with cognition, some features are uniquely associated with the general exposome factor, and some variance is shared between both.

## DISCUSSION

Neurodevelopment does not take place in a vacuum, but enmeshed in a complex, multifaceted web of environmental exposures and experiences that continually shape us over time. Understanding how individual differences in cognition emerge during childhood therefore requires the characterization of both an individual’s unique environment and their unique patterns of functional brain network organization. To characterize reproducible cross-sectional and longitudinal environment-brain-behavior associations, we conducted preregistered analyses using data from the ABCD Study^Ⓡ^ across two large-scale matched samples of youth. Using three previously validated dimensionality reduction approaches, we comprehensively measured and characterized the many inter-connected features of an individual’s exposome, mapped individual-specific patterns of functional brain organization, and trained cross-validated models to predict cognitive performance from held-out data. Our results highlight the substantial contribution of a child’s exposome to both functional brain network organization and cognitive development. In particular, we found that inter-individual differences in the exposome help to explain individual differences in cognition in children and predict cognitive abilities two years later over and above cognition at baseline. Associations between exposome scores and cognition remained significant over and above associations with socio-economic status and psychiatric medication use, and were highly consistent across sub-samples stratified by biological sex or racial identification. Moreover, we found that the exposome is reflected in children’s unique patterns of functional network topography, shedding light on both shared and unique contributions of the exposome and brain organization in shaping cognitive development. Together, these results bolster our understanding of how a child’s multidimensional environment may shape their unique patterns of brain network organization and cognitive functioning.

Understanding how individual differences in cognitive functioning emerge during childhood is a critical prerequisite for efforts that seek to promote healthy neurocognitive development. Individual differences in cognition that are observed during childhood are associated with academic performance (Cortés et al., 2019) and quality of life in youth (Klassen et al., 2004), as well as social, physical and mental health outcomes in adulthood (Agha et al., 2019; Richards et al., 2004; Moffitt et al., 2011). Moreover, cognitive deficits during childhood and adolescence are associated with heightened risk for psychopathology (Shanmugan et al., 2016), risk-taking behaviors (Shamosh et al., 2008), cardiovascular disease (Gow et al., 2011; Hart et al., 2004; Wraw et al., 2015) and all-cause mortality (Batty et al., 2007; Calvin et al., 2011). Our work provides the first longitudinal characterization of the relationships among a child’s exposome, personalized functional brain network topography and cognition in a large-scale dataset of youth. This work therefore lays the groundwork for future studies to causally test environmental interventions aimed at promoting healthy cognition and neurodevelopment in youth.

Ever since the “exposome” framework was put forward as a way to measure an individual’s multidimensional environment and experiences (Rappaport, 2011; Wild, 2005), numerous studies have uncovered associations between the exposome and a variety of outcomes, particularly physical health outcomes in adults (Vermeulen et al., 2020). The present study builds on recent work which has expanded this focus to include mental health outcomes in children (Barzilay et al., 2022; Moore et al., 2022; Pries et al., 2022). We have recently described in a cross sectional analysis that a single measure capturing the exposome explains ∼40% of variance in overall psychopathology at early adolescence (Moore et al., 2022). In the present study, we extend this work to investigate the longitudinal associations between the exposome and cognitive functioning in youth and demonstrate that the exposome also explains substantially more variance in cognition than measures derived from functional neuroimaging. When we compared predictive models trained on just a single variable (capturing a child’s general exposome) with models trained on a wealth of robust, personalized neuroimaging variables (the full pattern of an individual’s personalized functional brain network topography across seventeen large-scale networks and nearly sixty thousand vertices), we found that the single-variable exposome models yielded predictions that were *just as accurate or even better* than the brain network models. Given that cognitive impairments are among the most prevalent and least well understood symptoms of transdiagnostic psychiatric illness (Cisler and Koster, 2010; Cotrena et al., 2016; Gotlib and Joormann, 2010; Keller et al., 2022a, 2019; Russman Block et al., 2020), our results represent an important complement to existing findings relating children’s exposome to psychopathology.

Our observation that the exposome is reflected in patterns of functional brain network topography supports the idea that a child’s environment leaves a mark on their neurodevelopment. A plethora of prior work has characterized in detail how specific types of experiences or attributes of the environment may affect brain network structure and function (Miguel et al., 2019). Many of these studies have focused on the effects of adverse childhood experiences such as childhood maltreatment (Teicher et al., 2016) or socio-economic status (Leijser et al., 2018). While our study does not uncover causal associations between specific attributes of the environment and specific attributes of functional brain network organization, our observation that a general exposome factor was strongly and reproducibly associated with general cognitive functioning lends credence to the approach of quantifying multiple environmental features at once to capture potentially important additive or interactive effects. Furthermore, our findings provide validation for functional network topography as a potentially useful biomarker of environmental impacts on neurodevelopment. Future studies with wider longitudinal timepoints may further characterize the full extent and duration to which the mark of the exposome remains imprinted on functional network topography, as well as further delineating the impact that specific exposome sub-factors may have in shaping particular domains of cognition.

Moreover, burgeoning work in the field has highlighted the importance of the environment in shaping the *pace* of brain development (Tooley et al., 2021), with observations of heterochronous windows of developmental plasticity across different parts of the brain (Sydnor et al., 2022). Adolescence appears to be a particularly crucial sensitive period for the development of higher-order cortices (Larsen and Luna, 2018; Sisk and Gee, 2022), including prominently the functional brain networks that support cognition. Our observation that multivariate patterns of functional topography in association and attention networks specifically were most strongly related to exposome scores provides further evidence that these networks are in a sensitive period during this time: a period in which they are highly plastic, and therefore strongly influenced by the environment. Indeed, these potential sensitive windows for higher-order cognitive development may occur opportunistically during a period of life when it is most crucial to learn to adapt to the environment. For example, as attention networks continue to become refined throughout childhood and adolescence, varying types of environmental adversity may shape the development of these networks to optimize their function for allocating attention adaptively (e.g., heightening broad awareness to increase fast orientation to potential threats versus honing sharp focus amid distractions). Further understanding the precise spatial and temporal windows for sensitive periods across the brain and the influence of specific types of adversity may help future work to target interventions for maximal impact on healthy cognitive development.

An ongoing challenge in the field has been to strike an effective balance between specificity, accuracy, and personalization on the one hand and reproducibility, parsimony, and generalizability on the other hand. While approaches favoring specificity allow us to deeply characterize associations between the environment and neurodevelopment with increased granularity and accuracy at the level of the individual, approaches favoring reproducibility allow our findings to be more readily understandable and generalizable to the wider population. In the field of statistical learning, this conundrum is referred to as a tradeoff between model flexibility and model interpretability (James et al., 2023), with more complex models yielding a more accurate fit to training data at the risk of overfitting and not generalizing to new samples. In practice, this tradeoff plays out in our choice of study design: following the reproducibility crisis in psychology and neuroscience (Ioannidis, 2005), the field has shifted its preference toward large-scale studies that confer generalizability, sacrificing some degree of granularity as not all measures (e.g. of environment or cognition) can be included in all large-scale studies. In the present study, we have attempted to balance this tradeoff in several ways. First, we used as much detail as possible to comprehensively measure each child’s exposome across diverse environmental features, took a personalized neuroscience approach to define functional brain network organization at the level of the individual, and assessed multiple domains of cognitive functioning. Second, to counterbalance this level of detail, we used dimensionality reduction approaches for all three measurement modalities to derive more broad, generalizable descriptors of environment, brain and behavior. Third, we maximized our chances of reproducibility by pre-registering our analyses and hypotheses, using a rigorous cross-validation procedure to train and test our models, and choosing model-selection statistics (AIC and BIC) that balance the tradeoff between model flexibility and interpretability. This approach may be applied in future studies, including future longitudinal data releases from the ABCD Study^Ⓡ^.

### Limitations and Future Directions

While the present study took a rigorous approach to understanding associations among exposome, functional brain organization and cognition, our results should be considered in light of several limitations. First, it is challenging to comprehensively capture every possible aspect of a child’s complex, multidimensional environment and experiences. Our approach to defining the exposome longitudinally made use of data that were collected as part of the ABCD Study^Ⓡ^ and which were available at multiple time points. Although this meant we could make use of a large number of variables capturing a variety of features of each child’s environment, not all aspects of a child’s environment were captured and some aspects of the environment may have been better assessed than others. In particular, it is important to note that the variables available for our study across longitudinal timepoints did not address overt or covert racial discrimination, which have known effects on child and adolescent development (Trent et al., 2019). Future studies may therefore investigate a wider array of assessments to more fully capture each child’s complex, multidimensional environment. Second, as the ABCD Study^Ⓡ^ is observational, we cannot infer causality from exposome to brain/cognition. Nonetheless, the longitudinal design and the fact that the exposome at ages 9-10 contributes to cognition two years later supports the notion of directionality from exposome to cognition. Future work may also leverage methods for causal inference to address the question of causality (Shi and Norgeot, 2022; Wold, 1956), which is critical to inform preventive interventions on modifiable environmental exposures that can help promote healthy neurocognitive development

## Conclusion

Our results highlight the critical contributions of a child’s environment to functional brain network organization and cognitive development. By reproducibly characterizing the link between early childhood environments, personalized functional brain network organization, and individual differences in cognition in youth, this study advances our understanding of how our unique environments shape our individual minds and brains. This study also provides a framework by which to identify replicable environment-brain-behavior associations using a personalized neuroscience approach, laying a strong foundation for future studies to further investigate environmental influences on individual-specific trajectories of neurocognitive development.

## MATERIALS AND METHODS

### Participants

Data were drawn from the Adolescent Brain Cognitive Development℠ (ABCD) study (Volkow et al., 2018) baseline sample from the ABCD BIDS Community Collection (ABCC, ABCD-3165) (Feczko et al., 2021), which included *n*=11,878 children aged 9-10 years old and their parents/guardians collected across 21 sites. Inclusion criteria for this study included being within the desired age range (9-10 years old), English language proficiency in the children, and having the ability to provide informed consent (parent) and assent (child). Exclusion criteria included the presence of severe sensory, intellectual, medical or neurological issues that would have impacted the child’s ability to comply with the study protocol, as well as MRI scanner contraindications. We additionally excluded participants with incomplete data or excessive head motion, as described below.

To test the generalizability of our results, we repeated each of our analyses in both a discovery sample (*n*=5,139) and a separate replication sample (*n*=5,137) that were matched across multiple socio-demographic variables including age, sex, site, ethnicity, parent education, combined family income, and others (Cordova et al., 2021; Feczko et al., 2021). Socio-demographic characteristics of participants in the discovery and replication samples may be found in **Supplementary Table 2**. We observed no significant differences between participants in the discovery and replication samples across any socio-demographic variables.

### Cognitive Assessment

Participants completed a battery of cognitive assessments, including seven tasks from the NIH Toolbox (Picture Vocabulary, Flanker Test, List Sort Working Memory Task, Dimensional Change Card Sort Task, Pattern Comparison Processing Speed Task, Picture Sequence Memory Task, and the Oral Reading Test) (Weintraub et al., 2013) as well as two additional tasks (the Little Man Task and the Rey Auditory Verbal Learning Task) (Luciana et al., 2018). To reduce the dimensionality of these measures and focus our analyses on cognitive domains that explained the majority of behavioral variance in these tasks, we used scores in three previously-established cognitive domains derived from a prior study in this same dataset (Thompson et al., 2019): 1) general cognition, 2) executive function, and 3) learning/memory. In this study, a three-factor Bayesian Probabilistic Principal Components Analysis (BPPCA) model was applied to the aforementioned battery of nine cognitive tasks. Scores generated by varimax rotated loadings for this three-factor model for general cognition (highest loadings: Oral Reading Test, Picture Vocabulary, and Little Man Task), executive function (highest loadings: Pattern Comparison Processing Speed Task, Flanker Test, and Dimensional Change Card Sort Task), and learning/memory (highest loadings: Picture Sequence Memory Task, Rey Auditory and Verbal Learning Task, and List Sort Working Memory Task) were downloaded directly from the ABCD Data Exploration and Analysis Portal (DEAP). Given that not all of these nine cognitive tasks were collected at all longitudinal timepoints, it was not possible to use the same factor scores for our longitudinal (year two) analyses. Therefore, for all longitudinal linear mixed effects analyses, we instead used scores on five individual cognitive tasks that were assessed at both baseline and two-year time points: Picture Vocabulary, Flanker, Picture Sequence Memory, Pattern Comparison and Reading Recognition.

### Definition of Exposome Factors

Our aim was to define both general and specific exposome factors capturing a child’s unique, complex, multidimensional experiences and environment. To do so, we applied the same approach as in our previous cross-sectional work (Moore et al., 2022) to data collected across multiple longitudinal time points. We began with a large number of variables (354) assessing various aspects of a child’s environment and experiences. These variables were in multiple formats (continuous, ordinal, and nominal), different lengths (scales used in the ABCD Study^Ⓡ^ ranged from 2 to 59 items in length), and multiple sources (youth-report, parent-report, geo-coded data, etc.). The first step was to determine whether each scale should be reduced to data-driven summary scores rather than using individual items. This was determined largely by the relative lengths of the scales, where the goal was to avoid having longer scales (those with more items) or variable sets (e.g. 91 geographic variables) “dominate” the exposome factors in the final analysis (see below). Scales were also reduced to a summary score if their inter-item correlations were high and therefore likely to form a single cluster in the final analysis; for example, the three-item neighborhood safety scale was reduced to a single score for this reason.

Ultimately, twelve scales were reduced in this data-driven manner: youth-report School Risk and Protective Factors Survey (four sub-scores), youth-report Family Environment Scale (two sub-scores), parent-report Family Environment Scale (two sub-scores), youth-report Parent Monitoring Survey (one sub-score), parent-report Neighborhood Safety/Crime Survey (one sub-score), parent-report Community Risk and Protective Factors (one sub-score), parent-report Mexican-American Cultural Values Scale (four sub-scores), parent-report Parental Rules on Substance Use (one sub-score), parent-report Sports and Activities Involvement Questionnaire (three sub-scores), Traumatic Brain Injury sum scores (one sub-score), youth-report Youth Substance Use Attitudes Questionnaire (one sub-score), and youth-reported hours of screen time on various devices (one sub-score). Additionally, the address-/geocode-based measures of the neighborhood and state environment were reduced to eight sub-scores. The above “pre-reduction” steps were conducted using exploratory factor analysis (EFA), where the number of factors to extract was determined by a combination of interpretability and subjective evaluation of the scree plot. **Supplementary Table 8** shows the sub-scales resulting from the above analyses, along with the items composing them. Note that there are 30 sub-scales, while only 29 were used; sub-scale “dry_heat” was dropped from analyses because of difficulty in interpretation. Temperature was nonetheless accounted for by the “traditional_south_and_midwest” sub-score, which included a count of the number of “extreme heat days” experienced in a given year. Analyses in this “pre-reduction” step were conducted using the *psych* package (Revelle, 2019) in R. Note that, in addition to the 29 sub-scales created in this step, the final analysis (below) included parent education, household income, and a binary variable indicating whether the child’s parents were married, for a total of 32 variables.

For the final analysis of 32 variables we used an exploratory structural equation model (ESEM) (Asparouhov and Muthén, 2009) using bifactor rotation (BI-CF-QUARTIMAX) accounting for clustering by families, stratified by site, and constraining factor loadings to be equal across time points (ensuring longitudinal measurement invariance). The number of factors to extract was determined by a combination of interpretability and model fit, where fit was assessed using the comparative fit index (CFI; >0.90 acceptable), root mean-square error of approximation (RSMEA; <0.08 acceptable), and standardized root mean-square residual (SRMR; <0.08 acceptable) (Hu and Bentler, 1999). Analyses were conducted using Mplus version 8.4 (Muthén and Muthén, 2017).

**Supplementary Table 1** shows the results of the ESEM with one general factor and six orthogonal sub-factors using 32 exposome variables. Fit of the model is acceptable, with CFI = 0.94, RMSEA = 0.026 ± 0.001, and SRMR = 0.032. The general factor reflects the overall exposome (somewhat analogous to a first principal component), with the strongest indicators relating to socioeconomic status (household income = 0.780; neighborhood poverty = −0.695; parent education = 0.680) and weakest indicator related to neighborhood characteristics associated with retirement (−0.009). The first specific factor comprises school involvement, enjoyment, and performance, as well as some weaker influences of parental monitoring and youth-reported family turmoil. The second specific factor comprises all sub-scales of the Mexican American Cultural Values Scale (MACVS), which captures many aspects of family values, centrality, and culture. The third specific factor relates to family turmoil from the points-of-view of both the parents and youth, as well as a weak relation to substance abuse risk in that area. The fourth specific factor captures several aspects of the youth’s neighborhood, especially poverty, density, safety, and pollution. The fifth specific factor comprises extracurricular activities and traumatic brain injuries (TBIs) (possibly related). Finally, the sixth specific factor relates to screen time and (weakly) to peer deviance.

In addition to the model fit indices described above, bifactor models have specific reliability indices useful for evaluating the relative strengths of the factors, appropriateness and reliability of scores, etc. (Rodriguez et al., 2016). These indices are shown in **Supplementary Table 9**. Thorough discussion of these bifactor-specific metrics is beyond the present scope, but the most important for the present purposes is factor determinacy (Grice, 2001). Determinacy is an indication of how representative factor scores are of the factors from which they were derived. For example, note that the inter-factor correlations in **Supplementary** Figure 1 are not exactly 0 despite the factors being modeled as orthogonal; this is due to slight indeterminacy of the factors, and is always seen when scores are calculated from factors. The key value for our present purposes is the Factor Determinacy for the general exposome factor in **Supplementary Table 9**, which is 0.89. This value is beyond the conventionally used minimum of 0.80 recommended for score determinacy, suggesting the general factor score used in the present study is sufficiently determined.

### Image Processing

Imaging acquisition for the ABCD Study® has been described elsewhere (Casey et al., 2018). As previously described (Feczko et al., 2021), the ABCD-BIDS Community Collection (ABCC; Collection 3165) from which we drew our data was processed according to the ABCD-BIDS pipeline. This pipeline includes distortion correction and alignment, denoising with Advanced Normalization Tools (ANTS^88^), FreeSurfer^89^ segmentation, surface registration, and volume registration using FSL FLIRT rigid-body transformation.^90,91^ Processing was done according to the DCAN BOLD Processing (DBP) pipeline which included the following steps: 1) de-meaning and de-trending of all fMRI data with respect to time; 2) denoising using a general linear model with regressors for signal and movement; 3) bandpass filtering between 0.008 and 0.09 Hz using a 2nd order Butterworth filter; 4) applying the DBP respiratory motion filter (18.582 to 25.726 breaths per minute), and 5) applying DBP motion censoring (frames exceeding an FD threshold of 0.2mm or failing to pass outlier detection at +/- 3 standard deviations were discarded). Following preprocessing, we concatenated the time series data for both resting-state scans and three task-based scans (Monetary Incentive Delay Task, Stop-Signal Task, and Emotional N-Back Task) as in prior work (Cui et al., 2020) to maximize the available data for our analyses. Participants with fewer than 600 remaining TRs after motion censoring or who failed to pass ABCD quality control for their T1 or resting-state fMRI scan were excluded.

### Regularized Non-Negative Matrix Factorization

As previously described (Cui et al., 2020; Keller et al., 2022b; Li et al., 2017), we used non-negative matrix factorization (NMF) (Lee and Seung, 1999) to derive individualized functional networks. NMF identifies networks by positively weighting connectivity patterns that covary, leading to a highly specific and reproducible parts-based representation (Lee and Seung, 1999). Our approach was enhanced by a group consensus regularization term derived from previous work in an independent dataset (Cui et al., 2020) that preserves the inter-individual correspondence, as well as a data locality regularization term that makes the decomposition robust to imaging noise, improves spatial smoothness, and enhances functional coherence of the subject-specific functional networks (see Li et al. (2017) for details of the method; see also: https://github.com/hmlicas/Collaborative_Brain_Decomposition). As NMF requires nonnegative input, we re-scaled the data by shifting time courses of each vertex linearly to ensure all values were positive.^25^ As in prior work, to avoid features in greater numeric ranges dominating those in smaller numeric range, we further normalized the time course by its maximum value so that all the time points have values in the range of [0, 1]. For this study, we used identical parameter settings as in prior validation studies (Cui et al., 2020; Keller et al., 2022b; Li et al., 2017).

### Defining individualized networks

To facilitate group-level interpretations of individually-defined PFNs, we used a group consensus atlas from a previously published study in an independent dataset (Cui et al., 2020) as an initialization for individualized network definition. In this way, we also ensured spatial correspondence across all subjects. Details regarding the derivation of this group consensus atlas can be found in previous work (Cui et al., 2020). Briefly, group-level decomposition was performed multiple times on a subset of randomly selected subjects and the resulting decomposition results were fused to obtain one robust initialization that is highly reproducible. Next, inter-network similarity was calculated and normalized-cuts (Cai et al., 2011) based spectral clustering method was applied to group the PFNs into 17 clusters. For each cluster, the PFN with the highest overall similarity with all other PFNs within the same cluster was selected as the most representative. The resulting group-level network loading matrix *V* was transformed from *fsaverage5* space to *fslr* space using Connectome Workbench (Marcus et al., 2011), and thus the resultant matrix had 17 rows and 59,412 columns. Each row of this matrix represents a functional network, while each column represents the loadings of a given cortical vertex.

Using the previously-derived group consensus atlas (Cui et al., 2020) as a prior to ensure inter-individual correspondence, we derived each individual’s specific network atlas using NMF based on the acquired group networks (17 x 59,412 loading matrix) as initialization and each individual’s specific fMRI times series. See Li et al. (2017) for optimization details. This procedure yielded a loading matrix *V* (17 x 59,412 matrix) for each participant, where each row is a PFN, each column is a vertex, and the value quantifies the extent each vertex belongs to each network. This probabilistic (soft) definition can be converted into discrete (hard) network definitions for display and comparison with other methods (Kong et al., 2019; Wang et al., 2015; Yeo et al., 2011) by labeling each vertex according to its highest loading. Split-half reliability of the PFN loadings was found to be high in prior work (Keller et al., 2022b), indicating excellent reliability of this measure.

### Linear Mixed-Effects Models

We used linear mixed effects models (implemented with the “lme4” package in *R*) to assess associations between exposome factors and cognitive performance while accounting for both fixed and random predictors. All linear mixed effects models included fixed effects parameters for age, biological sex, site, as well as a random intercept for family (accounting for siblings). Sensitivity analyses were conducted to test whether results held with the additional inclusion of covariates for commonly-used measures of socio-economic status (household income and parental education) or psychiatric medication use (ADHD medications, Antidepressants, or Antipsychotics). We note that medication use was assessed using the PhenX instrument and coded as in our previous work (Shoval et al., 2021). Additionally, we conducted stratified analyses to ensure that our results held across categorical definitions of biological sex and self-reported racial identification.

### Ridge Regression Models

To uncover associations between the full multivariate pattern of PFN functional topography and each participant’s general exposome factor score, we trained linear ridge regression models using nested two-fold cross validation as in our previous work (Keller et al., 2022; Cui et al., 2020). In line with the recommendation that predictive models of brain-behavior associations be trained on multivariate patterns rather than univariate measures (Rosenberg and Finn, 2022), these predictive models were trained on concatenated network loading matrices across the 17 PFNs. Independent network models were also trained on loadings from specific networks. All models included covariates for age, sex, site, and motion (mean FD) that were regressed out separately in the training and testing sets prior to training the ridge regression models. These ridge regression models were trained and tested in our matched discovery and replication samples (Cordova et al., 2021; Feczko et al., 2021) using nested two-fold cross-validation (2F-CV), with outer 2F-CV estimating the generalizability of the model and the inner 2F-CV determining the optimal tuning parameter (λ) for the ridge regression. For the inner 2F-CV, one subset was selected to train the model under a given λ value in the range [1, 10, 100, 500, 1000, 5000, 10000, 15000, 20000] (Keller et al., 2022), and the remaining subset was used to test the model. This procedure was repeated 2 times such that each subset was used once as the testing dataset, resulting in two inner 2F-CV loops in total. For each λ value, the correlation *r* between the observed and predicted outcome as well as the mean absolute error (MAE) were calculated for each inner 2F-CV loop, and then averaged across the two inner loops. The sum of the mean correlation *r* and reciprocal of the mean MAE was defined as the inner prediction accuracy, and the λ with the highest inner prediction accuracy was chosen as the optimal λ (Keller et al., 2022; Cui et al., 2020).

To ensure that our matched discovery and replication sample selection procedure did not bias our results, we performed repeated random cross-validation over 100 iterations, each time randomly splitting the sample and repeating the nested 2F-CV procedure to generate a distribution of prediction accuracies for each model. Furthermore, we used permutation testing to generate null distributions for both the primary models and the repeated random cross-validation models by randomly shuffling the outcome variable. To ensure that our results were not overfit as a result of leakage across samples by the general exposome factor outcome variables derived in the whole sample, we also trained ridge regression models with the general exposome factor derived by two independent longitudinal bifactor analyses in the discovery and replication samples separately. Repeating our main analyses with these new predictive models, we find nearly identical results as shown in **Supplementary** Figure 3.

To investigate associations among the exposome, PFN topography, and cognition, we trained three additional ridge regression models using the same procedure as above: Model 1 (“Exp-Factor”) used only a participant’s general exposome score; Model 2 (“PFN Topography”), reported in our prior work (Keller et al., 2022b), used each participant’s multivariate pattern of personalized functional network topography; and Model 3 (“Exp-Factor + PFN Topography”) used a combination of the features in the first and second model types, hypothesized to perform best by capitalizing on both shared and unique variance across features. By evaluating model performance and comparing performance among the three model types, we were able to determine the relative contributions of a child’s exposome and functional brain network organization to cognitive functioning. Note that the model training procedure with ridge regression remained the same even for the single-variable model to allow the slope of the association between the general exposome factor and cognition to vary with the value of the ridge constraint during nested cross-validation for parameter tuning. This procedure was again repeated for our longitudinal prediction analysis by training these same models to predict cognitive performance on five tasks (Picture Vocabulary, Flanker, Picture Sequence Memory, Pattern Comparison and Reading Recognition) measured two years later when children were 11-12 years old. These models included additional covariates for baseline cognitive performance on each task. In all cases, models were compared by assessing the Akaike Information Criterion (AIC) (Akaike, 1998) and Bayesian Information Criterion (BIC) (Schwarz, 1978), which (unlike other measures like R^2^) take into account the number of features that the model is trained on, penalizing more complex models.

## Acknowledgment

This study was supported by grants from the National Institutes of Health: R01MH113550 (TDS), R01MH120482 (TDS), R01EB022573 (TDS & YF), R01MH123563 (RTS & TDS), R37MH125829 (TDS & DAF), R01MH123550 (RTS), R01MH112847 (RTS), K23MH120437 (RB). ASK was supported by a Neuroengineering and Medicine T32 Fellowship from the National Institute of Neurological Disorders and Stroke (5T32NS091006-08) and a Neurodevelopment and Psychosis T32 Fellowship from the National Institute of Mental Health (5T32MH019112-32). AP was supported by the Stanford School of Medicine Dean’s Fellowship. Additional support was provided by the Penn-CHOP Lifespan Brain Institute.

Data used in the preparation of this article were obtained from the Adolescent Brain Cognitive Development^SM^ (ABCD) Study (https://abcdstudy.org), held in the NIMH Data Archive (NDA). This is a multisite, longitudinal study designed to recruit more than 10,000 children age 9-10 and follow them over 10 years into early adulthood. The ABCD Study® is supported by the National Institutes of Health and additional federal partners under award numbers U01DA041048, U01DA050989, U01DA051016, U01DA041022, U01DA051018, U01DA051037, U01DA050987, U01DA041174, U01DA041106, U01DA041117, U01DA041028, U01DA041134, U01DA050988, U01DA051039, U01DA041156, U01DA041025, U01DA041120, U01DA051038, U01DA041148, U01DA041093, U01DA041089, U24DA041123, U24DA041147. A full list of supporters is available at https://abcdstudy.org/federal-partners.html. A listing of participating sites and a complete listing of the study investigators can be found at https://abcdstudy.org/consortium_members/. ABCD consortium investigators designed and implemented the study and/or provided data but did not necessarily participate in the analysis or writing of this report. This manuscript reflects the views of the authors and may not reflect the opinions or views of the NIH or ABCD consortium investigators. The ABCD data repository grows and changes over time. The ABCD data used in this report came from [NIMH Data Archive Digital Object Identifier 10.15154/1523041]. DOIs can be found at https://nda.nih.gov/abcd.

