## Supplementary Material for "A general exposome factor explains individual differences in functional brain network topography and cognition in youth"

| Variable | Exp-Factor | School | Family Values | Family Turmoil | Dense Urban Poverty | Extracurriculars | Screen Time |
|---|---|---|---|---|---|---|---|
| SRPF: Feel Involved in School | 0.075 | 0.641 | 0.012 | 0.023 | 0.003 | -0.043 | 0.065 |
| SRPF: Good Grades | -0.063 | 0.628 | 0.037 | -0.010 | -0.009 | -0.019 | -0.042 |
| SRPF: Enjoy School | 0.090 | 0.625 | 0.010 | -0.034 | 0.016 | 0.050 | -0.060 |
| SRPF: Positive Feedback in School | -0.104 | 0.581 | 0.019 | -0.002 | -0.010 | 0.072 | 0.018 |
| Parental Monitoring | 0.208 | 0.435 | 0.039 | -0.069 | 0.037 | -0.068 | 0.014 |
| FES: Family Non-Conflict - Youth Report | -0.182 | -0.278 | -0.034 | 0.259 | -0.068 | 0.035 | 0.033 |
| MACV: Family Image | **-0.437** | 0.032 | 0.739 | 0.001 | 0.040 | -0.010 | 0.018 |
| MACV: Caring/Security | -0.147 | 0.017 | 0.737 | -0.029 | -0.027 | 0.024 | -0.015 |
| MACV: Religiosity | **-0.465** | 0.046 | 0.397 | -0.011 | -0.141 | -0.014 | -0.047 |
| MACV: Independence | -0.247 | 0.000 | 0.385 | 0.030 | 0.111 | -0.007 | 0.020 |
| Parental Rules on Substance Use | 0.201 | -0.021 | -0.129 | 0.024 | 0.109 | 0.022 | 0.017 |
| FES: Family Conflict - Parent Report | -0.180 | -0.019 | -0.008 | 0.551 | 0.013 | 0.019 | -0.008 |
| FES: Family Non-Conflict - Parent Report | 0.012 | -0.030 | -0.071 | 0.474 | 0.025 | 0.023 | 0.012 |
| FES: Family Conflict - Youth Report | -0.271 | -0.257 | -0.014 | 0.347 | -0.150 | 0.086 | 0.011 |
| Substance Use Attitudes | 0.225 | -0.036 | -0.101 | 0.154 | 0.080 | -0.083 | 0.025 |
| ABGD: Dense/Suburban | -0.121 | 0.036 | 0.029 | 0.051 | 0.553 | 0.006 | -0.008 |
| ABGD: Crowding and Crime | **-0.311** | 0.016 | 0.100 | -0.085 | 0.499 | -0.047 | -0.050 |
| ABGD: Retirement and Group Living | -0.009 | -0.021 | -0.042 | -0.023 | 0.486 | 0.018 | 0.016 |
| ABGD: Poverty | **-0.695** | 0.012 | -0.026 | -0.022 | 0.393 | 0.007 | 0.010 |
| Neighborhood Safety | **0.398** | -0.016 | 0.122 | -0.085 | -0.352 | 0.025 | 0.022 |
| ABGD: Air Polution | -0.043 | 0.019 | -0.023 | 0.071 | 0.352 | -0.003 | 0.057 |
| ABGD: Traditional South and Midwest | **-0.415** | 0.040 | 0.035 | 0.018 | -0.293 | -0.079 | -0.020 |
| Parents Married | **0.572** | -0.002 | 0.059 | -0.013 | -0.264 | -0.064 | -0.130 |
| ABGD: Ozone | 0.122 | -0.049 | 0.011 | -0.066 | 0.253 | 0.024 | 0.037 |
| Household Income | **0.780** | -0.022 | 0.008 | 0.034 | -0.240 | 0.001 | -0.017 |
| Parental Education | **0.680** | -0.028 | -0.044 | 0.060 | -0.214 | 0.096 | -0.016 |
| SAIQ: Miscellaneous Sports/Activities | **0.507** | 0.004 | 0.032 | 0.065 | -0.004 | 0.433 | 0.018 |
| SAIQ: Arts and Individual Sports/Activities | **0.475** | 0.052 | 0.003 | 0.016 | 0.043 | 0.379 | -0.003 |
| SAIQ: Competitive/High-Impact Athletics | 0.223 | -0.008 | 0.047 | 0.084 | -0.085 | 0.337 | 0.021 |
| TBIs | 0.085 | -0.034 | -0.007 | 0.053 | -0.013 | 0.178 | -0.003 |
| Screen Time | -0.235 | -0.063 | -0.006 | 0.018 | -0.004 | 0.032 | 0.393 |
| Peer Deviance | -0.086 | -0.086 | -0.016 | 0.046 | -0.006 | 0.017 | 0.099 |

Youth Report
Parent Report
Geocoded Data

**Supplementary Table 1. Exposome Score Bifactor Loadings.** Factor loadings from the longitudinal exploratory bifactor analysis (see Methods section for details on the analysis approach). Shaded gray boxes indicate the variables that load most strongly onto each sub-factor score. *Abbreviations:* SRPF: School Risk and Protective Factors Survey; FES: Family Environment Scale - Family Conflict Subscale; MACV: Mexican American Cultural Values Scale; ABGD: Address-Based Geographic Data; SAIQ: Sports and Activities Involvement Questionnaire.

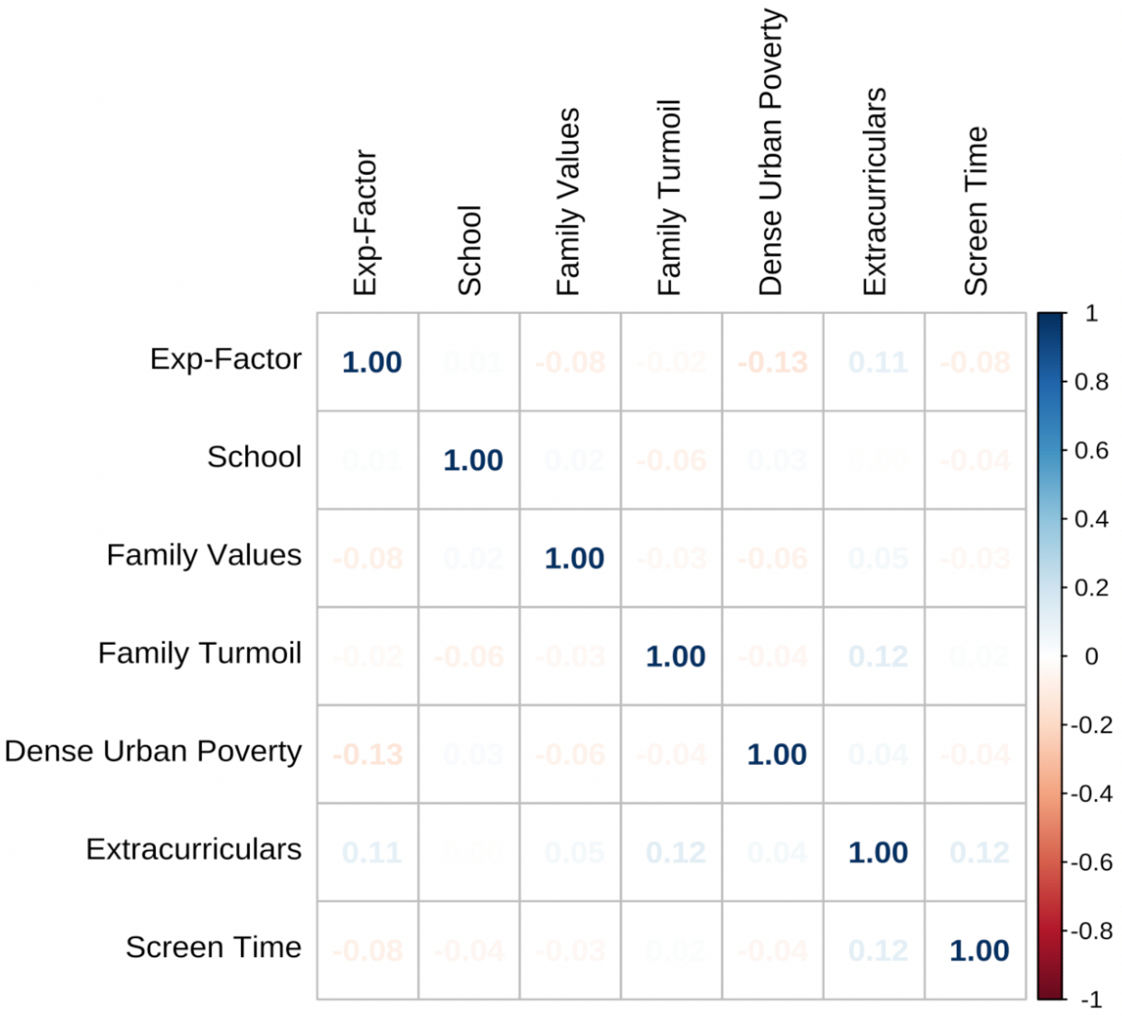


**Supplementary Figure 1. The general exposome factor (Exp-Factor) and specific exposome sub-factors are all orthogonal to one another.** Correlation matrix among all factors from the longitudinal bifactor analysis.

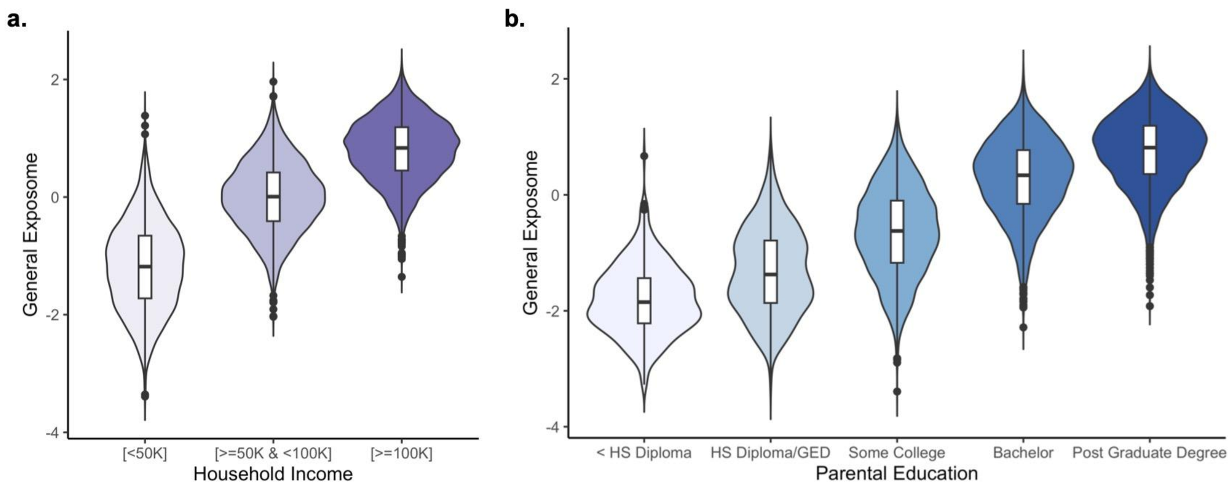


**Supplementary Figure 2. The general exposome factor is positively associated with measures of socio-economic status.** A one-way analysis of variance (ANOVA) confirmed that the general exposome factor varied significantly by both household income ($F(2,10528) = 12797.41$, $p < .001$) and parental education ($F(4,10528) = 919.91$, $p < .001$), with higher general exposome factor scores associated with higher socio-economic status.

| | Total | Discovery | Replication | P-value |
|---|---|---|---|---|
| Age (Months) | 119.0 (±7.5) | 119.0 (±7.5) | 119.0 (±7.5) | 0.80 |
| Sex | 4,914 (47.8%) | 2,493 (48.5%) | 2,421 (47.1%) | 0.16 |
| **Household Income** | | | | **0.74** |
| [<50K] | 2,970 (28.9%) | 1,470 (28.6%) | 1,500 (29.2%) | |
| [>=50K & <100K] | 2,932 (28.5%) | 1,481 (28.8%) | 1,451 (28.2%) | |
| [>=100K] | 4,374 (42.6%) | 2,188 (42.6%) | 2,186 (42.6%) | |
| **Race** | | | | **0.79** |
| White | 6,810 (66.3%) | 3,442 (67.0%) | 3,368 (65.6%) | |
| Black | 1,501 (14.6%) | 735 (14.3%) | 766 (14.9%) | |
| Asian | 227 (2.2%) | 112 (2.2%) | 115 (2.2%) | |
| AIAN/NHPI | 63 (0.6%) | 32 (0.6%) | 31 (0.6%) | |
| Other | 398 (3.9%) | 195 (3.8%) | 203 (4.0%) | |
| Mixed | 1,277 (12.4%) | 623 (12.1%) | 654 (12.7%) | |
| **Parent Education** | | | | **0.085** |
| < HS Diploma | 390 (3.8%) | 217 (4.2%) | 173 (3.4%) | |
| HS Diploma/GED | 860 (8.4%) | 413 (8.0%) | 447 (8.7%) | |
| Some College | 2,631 (25.6%) | 1,292 (25.1%) | 1,339 (26.1%) | |
| Bachelor | 2,714 (26.4%) | 1,383 (26.9%) | 1,331 (25.9%) | |
| Post Graduate Degree | 3,681 (35.8%) | 1,834 (35.7%) | 1,847 (36.0%) | |

**Supplementary Table 2. Demographic characteristics in the matched discovery (*n*=5,139) and replication (*n*=5,137) samples.** *Abbreviations*: AIAN = American Indian/Alaska Native; NHPI = Native Hawaiian and other Pacific Islander; HS = High School; GED = General Educational Development; CBCL = Child Behavior Checklist.

| | Picture Vocabulary | | | | Flanker | | | | Picture Sequence Memory | | | | Pattern Comparison | | | | Reading Recognition | | | |
|---|---|---|---|---|---|---|---|---|---|---|---|---|---|---|---|---|---|---|---|---|
| *Predictors* | $\beta$ | *Std. Error* | *t* | $p_{bonf}$ | $\beta$ | *Std. Error* | *t* | $p_{bonf}$ | $\beta$ | *Std. Error* | *t* | $p_{bonf}$ | $\beta$ | *Std. Error* | *t* | $p_{bonf}$ | $\beta$ | *Std. Error* | *t* | $p_{bonf}$ |
| **Discovery Sample** | | | | | | | | | | | | | | | | | | | | |
| Intercept | -0.02 | 0.04 | -0.65 | 1.00 | -0.06 | 0.03 | -1.81 | $7.08 \times 10^{-1}$ | -0.01 | 0.03 | -0.34 | 1.00 | 0.05 | 0.04 | 1.29 | 1.00 | -0.02 | 0.05 | -0.35 | 1.00 |
| Age | 0.23 | 0.01 | 19.88 | **$9.62 \times 10^{-84}$** | 0.18 | 0.01 | 13.22 | **$2.76 \times 10^{-38}$** | 0.11 | 0.01 | 8.25 | **$1.93 \times 10^{-15}$** | 0.22 | 0.01 | 16.48 | **$1.64 \times 10^{-58}$** | 0.21 | 0.01 | 16.80 | **$1.07 \times 10^{-60}$** |
| Sex | 0.07 | 0.02 | 2.81 | **$4.96 \times 10^{-2}$** | 0.05 | 0.03 | 1.87 | $6.21 \times 10^{-1}$ | -0.12 | 0.03 | -4.27 | **$1.99 \times 10^{-4}$** | -0.13 | 0.03 | -4.72 | **$2.40 \times 10^{-5}$** | -0.00 | 0.03 | -0.14 | 1.00 |
| Exp-Factor | 0.50 | 0.01 | 36.06 | **$4.95 \times 10^{-253}$** | 0.20 | 0.02 | 13.33 | **$7.06 \times 10^{-39}$** | 0.22 | 0.01 | 14.85 | **$6.82 \times 10^{-48}$** | 0.12 | 0.02 | 8.06 | **$9.72 \times 10^{-15}$** | 0.43 | 0.01 | 28.99 | **$2.90 \times 10^{-170}$** |
| School | -0.02 | 0.01 | -1.78 | $7.47 \times 10^{-1}$ | -0.01 | 0.01 | -0.57 | 1.00 | 0.02 | 0.01 | 1.06 | 1.00 | 0.03 | 0.01 | 2.03 | $4.27 \times 10^{-1}$ | -0.00 | 0.01 | -0.18 | 1.00 |
| Family Values | -0.10 | 0.01 | -7.06 | **$1.91 \times 10^{-11}$** | 0.02 | 0.02 | 1.49 | 1.00 | -0.03 | 0.02 | -2.10 | $3.60 \times 10^{-1}$ | 0.01 | 0.02 | 0.67 | 1.00 | -0.01 | 0.01 | -0.63 | 1.00 |
| Family Turmoil | 0.01 | 0.01 | 0.87 | 1.00 | 0.01 | 0.01 | 0.46 | 1.00 | 0.00 | 0.01 | 0.13 | 1.00 | -0.02 | 0.01 | -1.17 | 1.00 | 0.01 | 0.01 | 0.58 | 1.00 |
| Dense Urban Poverty | 0.01 | 0.01 | 0.49 | 1.00 | -0.01 | 0.01 | -0.97 | 1.00 | -0.01 | 0.01 | -0.47 | 1.00 | -0.04 | 0.02 | -2.40 | $1.64 \times 10^{-1}$ | 0.05 | 0.02 | 3.23 | **$1.25 \times 10^{-2}$** |
| Extracurriculars | -0.02 | 0.02 | -1.00 | 1.00 | 0.01 | 0.02 | 0.64 | 1.00 | 0.02 | 0.02 | 0.92 | 1.00 | -0.01 | 0.02 | -0.32 | 1.00 | -0.01 | 0.02 | -0.45 | 1.00 |
| Screen Time | -0.03 | 0.02 | -1.62 | 1.00 | -0.03 | 0.02 | -1.57 | 1.00 | -0.12 | 0.02 | -5.49 | **$4.26 \times 10^{-7}$** | -0.04 | 0.02 | -2.03 | $4.27 \times 10^{-1}$ | -0.08 | 0.02 | -3.82 | **$1.35 \times 10^{-3}$** |
| **Replication Sample** | | | | | | | | | | | | | | | | | | | | |
| Intercept | -0.05 | 0.03 | -1.37 | 1.00 | -0.04 | 0.04 | -1.10 | 1.00 | 0.01 | 0.03 | 0.19 | 1.00 | 0.03 | 0.04 | 0.90 | 1.00 | -0.02 | 0.05 | -0.44 | 1.00 |
| Age | 0.25 | 0.01 | 22.01 | **$1.06 \times 10^{-101}$** | 0.16 | 0.01 | 12.20 | **$9.41 \times 10^{-33}$** | 0.11 | 0.01 | 8.09 | **$7.69 \times 10^{-15}$** | 0.22 | 0.01 | 16.05 | **$1.39 \times 10^{-55}$** | 0.24 | 0.01 | 19.45 | **$2.22 \times 10^{-80}$** |
| Sex | 0.07 | 0.02 | 2.89 | **$3.87 \times 10^{-2}$** | 0.02 | 0.03 | 0.62 | 1.00 | -0.11 | 0.03 | -3.95 | **$7.99 \times 10^{-4}$** | -0.11 | 0.03 | -4.03 | **$5.62 \times 10^{-4}$** | 0.04 | 0.03 | 1.67 | $9.58 \times 10^{-1}$ |
| Exp-Factor | 0.48 | 0.01 | 36.31 | **$4.80 \times 10^{-256}$** | 0.22 | 0.02 | 14.76 | **$2.54 \times 10^{-47}$** | 0.22 | 0.01 | 14.78 | **$1.85 \times 10^{-47}$** | 0.15 | 0.02 | 9.62 | **$9.75 \times 10^{-21}$** | 0.43 | 0.01 | 29.58 | **$1.12 \times 10^{-176}$** |
| School | -0.02 | 0.01 | -1.25 | 1.00 | 0.00 | 0.01 | 0.23 | 1.00 | 0.01 | 0.01 | 0.76 | 1.00 | 0.06 | 0.01 | 4.03 | **$5.66 \times 10^{-4}$** | -0.01 | 0.01 | -1.09 | 1.00 |
| Family Values | -0.10 | 0.01 | -7.03 | **$2.26 \times 10^{-11}$** | -0.02 | 0.02 | -1.55 | 1.00 | -0.01 | 0.02 | -0.74 | 1.00 | -0.02 | 0.02 | -1.54 | 1.00 | -0.02 | 0.01 | -1.26 | 1.00 |
| Family Turmoil | -0.01 | 0.01 | -1.15 | 1.00 | -0.00 | 0.01 | -0.25 | 1.00 | 0.01 | 0.01 | 0.95 | 1.00 | -0.01 | 0.01 | -0.68 | 1.00 | -0.01 | 0.01 | -0.54 | 1.00 |
| Dense Urban Poverty | -0.00 | 0.01 | -0.32 | 1.00 | -0.02 | 0.02 | -1.53 | 1.00 | -0.04 | 0.01 | -2.70 | $6.94 \times 10^{-2}$ | 0.00 | 0.02 | 0.11 | 1.00 | 0.02 | 0.01 | 1.39 | 1.00 |
| Extracurriculars | -0.01 | 0.02 | -0.68 | 1.00 | 0.01 | 0.02 | 0.52 | 1.00 | 0.00 | 0.02 | 0.08 | 1.00 | -0.00 | 0.02 | -0.11 | 1.00 | -0.02 | 0.02 | -1.02 | 1.00 |
| Screen Time | -0.06 | 0.02 | -3.22 | **$1.28 \times 10^{-2}$** | -0.03 | 0.02 | -1.28 | 1.00 | -0.08 | 0.02 | -3.79 | **$1.50 \times 10^{-3}$** | -0.07 | 0.02 | -3.07 | **$2.13 \times 10^{-2}$** | -0.06 | 0.02 | -3.21 | **$1.33 \times 10^{-2}$** |

**Supplementary Table 3. Exp-Factor is significantly associated with cognition.** Across all five cognitive tasks and across both the discovery and replication sub-samples, the general exposome factor (Exp-Factor) is positively associated with cognition. These effects held with the inclusion of all six orthogonal exposome sub-factors as covariates and survived Bonferroni correction for multiple comparisons. Note that random effects for site and family are also included as covariates in these models.

| | Picture Vocabulary | | | | Flanker | | | | Picture Sequence Memory | | | | Pattern Comparison | | | | Reading Recognition | | | |
|---|---|---|---|---|---|---|---|---|---|---|---|---|---|---|---|---|---|---|---|---|
| *Predictors* | $\beta$ | *Std. Error* | *t* | $p_{bonf}$ | $\beta$ | *Std. Error* | *t* | $p_{bonf}$ | $\beta$ | *Std. Error* | *t* | $p_{bonf}$ | $\beta$ | *Std. Error* | *t* | $p_{bonf}$ | $\beta$ | *Std. Error* | *t* | $p_{bonf}$ |
| **Discovery Sample** | | | | | | | | | | | | | | | | | | | | |
| Intercept | -0.23 | 0.09 | -2.68 | $1.17 \times 10^{-1}$ | -0.18 | 0.10 | -1.84 | 1.00 | -0.09 | 0.10 | -0.99 | 1.00 | 0.19 | 0.10 | 1.95 | $8.14 \times 10^{-1}$ | -0.29 | 0.10 | -3.00 | **$4.31 \times 10^{-2}$** |
| Age | 0.23 | 0.01 | 20.08 | **$3.20 \times 10^{-85}$** | 0.18 | 0.01 | 13.28 | **$2.26 \times 10^{-38}$** | 0.11 | 0.01 | 8.35 | **$1.44 \times 10^{-15}$** | 0.22 | 0.01 | 16.46 | **$3.96 \times 10^{-58}$** | 0.21 | 0.01 | 17.04 | **$3.44 \times 10^{-62}$** |
| Sex | 0.07 | 0.02 | 2.81 | $7.89 \times 10^{-2}$ | 0.05 | 0.03 | 1.89 | $9.37 \times 10^{-1}$ | -0.12 | 0.03 | -4.30 | **$2.82 \times 10^{-4}$** | -0.13 | 0.03 | -4.67 | **$5.05 \times 10^{-5}$** | -0.00 | 0.03 | -0.16 | 1.00 |
| Household Income [50K-100K] | -0.05 | 0.04 | -1.06 | 1.00 | 0.06 | 0.05 | 1.19 | 1.00 | -0.06 | 0.05 | -1.31 | 1.00 | -0.07 | 0.05 | -1.52 | 1.00 | -0.01 | 0.05 | -0.14 | 1.00 |
| Household Income [>100K] | -0.13 | 0.05 | -2.35 | $3.02 \times 10^{-1}$ | 0.03 | 0.06 | 0.49 | 1.00 | -0.07 | 0.06 | -1.05 | 1.00 | -0.16 | 0.06 | -2.61 | $1.45 \times 10^{-1}$ | -0.09 | 0.06 | -1.54 | 1.00 |
| Parent Educ: HS Diploma/GED | 0.14 | 0.07 | 1.94 | $8.44 \times 10^{-1}$ | -0.02 | 0.08 | -0.26 | 1.00 | 0.13 | 0.08 | 1.48 | 1.00 | -0.07 | 0.08 | -0.80 | 1.00 | 0.18 | 0.08 | 2.25 | $3.97 \times 10^{-1}$ |
| Parent Educ: Some College | 0.25 | 0.07 | 3.59 | **$5.39 \times 10^{-3}$** | 0.10 | 0.08 | 1.24 | 1.00 | 0.08 | 0.08 | 0.99 | 1.00 | -0.06 | 0.08 | -0.80 | 1.00 | 0.27 | 0.08 | 3.63 | **$4.59 \times 10^{-3}$** |
| Parent Educ: Bachelor | 0.29 | 0.08 | 3.72 | **$3.25 \times 10^{-3}$** | 0.12 | 0.09 | 1.34 | 1.00 | 0.14 | 0.09 | 1.55 | 1.00 | -0.03 | 0.09 | -0.33 | 1.00 | 0.34 | 0.08 | 4.03 | **$8.91 \times 10^{-4}$** |
| Parent Educ: Post Graduate | 0.38 | 0.08 | 4.62 | **$6.18 \times 10^{-5}$** | 0.08 | 0.09 | 0.82 | 1.00 | 0.20 | 0.09 | 2.16 | $4.97 \times 10^{-1}$ | -0.04 | 0.09 | -0.43 | 1.00 | 0.42 | 0.09 | 4.82 | **$2.40 \times 10^{-5}$** |
| Exp-Factor | 0.47 | 0.03 | 17.51 | **$1.68 \times 10^{-65}$** | 0.18 | 0.03 | 5.86 | **$7.83 \times 10^{-8}$** | 0.20 | 0.03 | 6.89 | **$9.76 \times 10^{-11}$** | 0.17 | 0.03 | 5.71 | **$1.89 \times 10^{-7}$** | 0.39 | 0.03 | 13.25 | **$3.27 \times 10^{-38}$** |
| School | -0.02 | 0.01 | -1.58 | 1.00 | -0.01 | 0.01 | -0.38 | 1.00 | 0.01 | 0.01 | 1.02 | 1.00 | 0.03 | 0.01 | 1.95 | $8.13 \times 10^{-1}$ | 0.00 | 0.01 | 0.07 | 1.00 |
| Family Values | -0.10 | 0.01 | -6.87 | **$1.18 \times 10^{-10}$** | 0.02 | 0.02 | 1.36 | 1.00 | -0.03 | 0.02 | -1.94 | $8.34 \times 10^{-1}$ | 0.01 | 0.02 | 0.90 | 1.00 | -0.01 | 0.01 | -0.55 | 1.00 |
| Family Turmoil | 0.01 | 0.01 | 0.74 | 1.00 | 0.00 | 0.02 | 0.32 | 1.00 | 0.00 | 0.02 | 0.05 | 1.00 | -0.01 | 0.02 | -0.79 | 1.00 | 0.00 | 0.01 | 0.34 | 1.00 |
| Dense Urban Poverty | 0.01 | 0.01 | 0.46 | 1.00 | -0.01 | 0.02 | -0.43 | 1.00 | -0.01 | 0.01 | -0.48 | 1.00 | -0.04 | 0.02 | -2.80 | $8.15 \times 10^{-2}$ | 0.05 | 0.02 | 3.32 | **$1.44 \times 10^{-2}$** |
| Extracurriculars | -0.03 | 0.02 | -1.73 | 1.00 | 0.01 | 0.02 | 0.68 | 1.00 | 0.01 | 0.02 | 0.48 | 1.00 | -0.01 | 0.02 | -0.69 | 1.00 | -0.02 | 0.02 | -1.08 | 1.00 |
| Screen Time | -0.03 | 0.02 | -1.82 | 1.00 | -0.04 | 0.02 | -1.80 | 1.00 | -0.12 | 0.02 | -5.40 | **$1.10 \times 10^{-6}$** | -0.04 | 0.02 | -1.76 | 1.00 | -0.08 | 0.02 | -4.03 | **$8.90 \times 10^{-4}$** |
| **Replication Sample** | | | | | | | | | | | | | | | | | | | | |
| Intercept | -0.10 | 0.09 | -1.11 | 1.00 | -0.18 | 0.10 | -1.78 | 1.00 | 0.04 | 0.10 | 0.40 | 1.00 | 0.07 | 0.10 | 0.70 | 1.00 | -0.30 | 0.10 | -2.98 | **$4.62 \times 10^{-2}$** |
| Age | 0.25 | 0.01 | 22.16 | **$9.48 \times 10^{-103}$** | 0.17 | 0.01 | 12.26 | **$6.75 \times 10^{-33}$** | 0.11 | 0.01 | 8.07 | **$1.37 \times 10^{-14}$** | 0.22 | 0.01 | 16.07 | **$1.68 \times 10^{-55}$** | 0.24 | 0.01 | 19.61 | **$1.88 \times 10^{-81}$** |
| Sex | 0.07 | 0.02 | 3.08 | **$3.34 \times 10^{-2}$** | 0.02 | 0.03 | 0.70 | 1.00 | -0.11 | 0.03 | -3.95 | **$1.28 \times 10^{-3}$** | -0.11 | 0.03 | -3.96 | **$1.20 \times 10^{-3}$** | 0.04 | 0.03 | 1.75 | 1.00 |
| Household Income [50K-100K] | -0.11 | 0.04 | -2.62 | $1.41 \times 10^{-1}$ | -0.04 | 0.05 | -0.88 | 1.00 | 0.02 | 0.05 | 0.51 | 1.00 | -0.03 | 0.05 | -0.62 | 1.00 | -0.00 | 0.04 | -0.04 | 1.00 |
| Household Income [>100K] | -0.21 | 0.05 | -4.16 | **$5.28 \times 10^{-4}$** | -0.06 | 0.06 | -1.04 | 1.00 | 0.01 | 0.06 | 0.25 | 1.00 | -0.11 | 0.06 | -1.84 | 1.00 | -0.10 | 0.06 | -1.77 | 1.00 |
| Parent Educ: HS Diploma/GED | -0.01 | 0.08 | -0.14 | 1.00 | 0.02 | 0.09 | 0.25 | 1.00 | -0.14 | 0.09 | -1.61 | 1.00 | 0.04 | 0.09 | 0.44 | 1.00 | 0.12 | 0.08 | 1.48 | 1.00 |
| Parent Educ: Some College | 0.10 | 0.07 | 1.40 | 1.00 | 0.16 | 0.09 | 1.82 | 1.00 | -0.08 | 0.09 | -0.87 | 1.00 | 0.00 | 0.09 | 0.04 | 1.00 | 0.27 | 0.08 | 3.32 | **$1.43 \times 10^{-2}$** |
| Parent Educ: Bachelor | 0.24 | 0.08 | 2.98 | **$4.70 \times 10^{-2}$** | 0.24 | 0.09 | 2.53 | $1.85 \times 10^{-1}$ | -0.04 | 0.10 | -0.45 | 1.00 | 0.02 | 0.09 | 0.18 | 1.00 | 0.36 | 0.09 | 4.10 | **$6.59 \times 10^{-4}$** |
| Parent Educ: Post Graduate | 0.28 | 0.09 | 3.26 | **$1.78 \times 10^{-2}$** | 0.23 | 0.10 | 2.30 | $3.42 \times 10^{-1}$ | -0.00 | 0.10 | -0.05 | 1.00 | 0.04 | 0.10 | 0.37 | 1.00 | 0.43 | 0.09 | 4.70 | **$4.26 \times 10^{-5}$** |
| Exp-Factor | 0.48 | 0.03 | 18.56 | **$3.05 \times 10^{-73}$** | 0.19 | 0.03 | 6.59 | **$7.97 \times 10^{-10}$** | 0.19 | 0.03 | 6.53 | **$1.17 \times 10^{-9}$** | 0.18 | 0.03 | 5.95 | **$4.55 \times 10^{-8}$** | 0.38 | 0.03 | 13.47 | **$1.79 \times 10^{-39}$** |
| School | -0.01 | 0.01 | -1.13 | 1.00 | 0.01 | 0.01 | 0.39 | 1.00 | 0.01 | 0.01 | 0.81 | 1.00 | 0.06 | 0.01 | 3.98 | **$1.13 \times 10^{-3}$** | -0.01 | 0.01 | -0.87 | 1.00 |
| Family Values | -0.09 | 0.01 | -6.60 | **$7.30 \times 10^{-10}$** | -0.02 | 0.02 | -1.47 | 1.00 | -0.01 | 0.02 | -0.81 | 1.00 | -0.02 | 0.02 | -1.24 | 1.00 | -0.02 | 0.01 | -1.07 | 1.00 |
| Family Turmoil | -0.02 | 0.01 | -1.40 | 1.00 | -0.01 | 0.02 | -0.61 | 1.00 | 0.01 | 0.02 | 0.73 | 1.00 | -0.01 | 0.02 | -0.43 | 1.00 | -0.02 | 0.01 | -1.06 | 1.00 |
| Dense Urban Poverty | -0.01 | 0.01 | -0.52 | 1.00 | -0.02 | 0.02 | -1.17 | 1.00 | -0.04 | 0.02 | -2.40 | $2.64 \times 10^{-1}$ | -0.00 | 0.02 | -0.23 | 1.00 | 0.03 | 0.02 | 1.75 | 1.00 |
| Extracurriculars | -0.02 | 0.02 | -1.54 | 1.00 | 0.00 | 0.02 | 0.18 | 1.00 | 0.00 | 0.02 | 0.06 | 1.00 | -0.01 | 0.02 | -0.36 | 1.00 | -0.03 | 0.02 | -1.49 | 1.00 |
| Screen Time | -0.05 | 0.02 | -2.75 | $9.60 \times 10^{-2}$ | -0.03 | 0.02 | -1.24 | 1.00 | -0.08 | 0.02 | -3.63 | **$4.58 \times 10^{-3}$** | -0.06 | 0.02 | -2.83 | $7.38 \times 10^{-2}$ | -0.06 | 0.02 | -3.11 | **$2.96 \times 10^{-2}$** |

**Supplementary Table 4. Exp-Factor at baseline assessment is significantly associated with cognition over and above standard measures of socio-economic status.** Across all five cognitive tasks and across both the discovery and replication sub-samples, the general exposome factor (Exp-Factor) is positively associated with cognition over and above standard measures of socio-economic status (household income and parental education). Note that random effects for site and family are also included as covariates in these models and that the effects of interest also remained significant in longitudinal models associating the general exposome factor with cognitive performance at the two year follow-up..

| | Picture Vocabulary | | | | Flanker | | | | Picture Sequence Memory | | | | Pattern Comparison | | | | Reading Recognition | | | |
|---|---|---|---|---|---|---|---|---|---|---|---|---|---|---|---|---|---|---|---|---|
| *Predictors* | $\beta$ | *Std. Error* | *t* | $p_{bonf}$ | $\beta$ | *Std. Error* | *t* | $p_{bonf}$ | $\beta$ | *Std. Error* | *t* | $p_{bonf}$ | $\beta$ | *Std. Error* | *t* | $p_{bonf}$ | $\beta$ | *Std. Error* | *t* | $p_{bonf}$ |
| **Discovery Sample** | | | | | | | | | | | | | | | | | | | | |
| Intercept | -0.00 | 0.04 | -0.04 | 1.00 | -0.02 | 0.04 | -0.48 | 1.00 | 0.02 | 0.04 | 0.54 | 1.00 | 0.07 | 0.05 | 1.40 | 1.00 | 0.01 | 0.05 | 0.25 | 1.00 |
| Age | 0.23 | 0.02 | 15.13 | **$1.30 \times 10^{-48}$** | 0.17 | 0.02 | 9.58 | **$2.76 \times 10^{-20}$** | 0.11 | 0.02 | 6.18 | **$9.53 \times 10^{-19}$** | 0.22 | 0.02 | 11.99 | **$3.11 \times 10^{-31}$** | 0.20 | 0.02 | 12.33 | **$6.05 \times 10^{-33}$** |
| Sex | 0.09 | 0.03 | 2.88 | $5.17 \times 10^{-2}$ | 0.09 | 0.04 | 2.53 | $1.50 \times 10^{-1}$ | -0.09 | 0.04 | -2.33 | $2.62 \times 10^{-1}$ | -0.12 | 0.04 | -3.08 | **$2.75 \times 10^{-2}$** | 0.05 | 0.03 | 1.47 | 1.00 |
| ADHD Medications | -0.07 | 0.06 | -1.15 | 1.00 | -0.24 | 0.07 | -3.43 | **$7.87 \times 10^{-3}$** | -0.12 | 0.07 | -1.67 | 1.00 | -0.17 | 0.07 | -2.35 | $2.47 \times 10^{-1}$ | -0.16 | 0.07 | -2.38 | $2.24 \times 10^{-1}$ |
| Antidepressants | 0.17 | 0.11 | 1.53 | 1.00 | -0.02 | 0.13 | -0.17 | 1.00 | -0.01 | 0.14 | -0.10 | 1.00 | 0.01 | 0.13 | 0.08 | 1.00 | 0.05 | 0.12 | 0.45 | 1.00 |
| Antipsychotics | -0.14 | 0.22 | -0.65 | 1.00 | -0.52 | 0.25 | -2.04 | $5.43 \times 10^{-1}$ | -0.41 | 0.27 | -1.55 | 1.00 | 0.33 | 0.26 | 1.27 | 1.00 | -0.19 | 0.24 | -0.81 | 1.00 |
| Exp-Factor | 0.46 | 0.02 | 23.85 | **$2.46 \times 10^{-113}$** | 0.19 | 0.02 | 9.44 | **$1.00 \times 10^{-19}$** | 0.21 | 0.02 | 10.21 | **$6.10 \times 10^{-23}$** | 0.11 | 0.02 | 5.07 | **$5.55 \times 10^{-6}$** | 0.37 | 0.02 | 17.63 | **$6.54 \times 10^{-65}$** |
| School | -0.01 | 0.02 | -0.82 | 1.00 | -0.01 | 0.02 | -0.42 | 1.00 | 0.02 | 0.02 | 1.09 | 1.00 | 0.03 | 0.02 | 1.42 | 1.00 | -0.01 | 0.02 | -0.44 | 1.00 |
| Family Values | -0.12 | 0.02 | -6.49 | **$1.34 \times 10^{-9}$** | -0.01 | 0.02 | -0.65 | 1.00 | -0.03 | 0.02 | -1.27 | 1.00 | -0.02 | 0.02 | -0.91 | 1.00 | -0.06 | 0.02 | -2.91 | **$4.67 \times 10^{-2}$** |
| Family Turmoil | -0.00 | 0.02 | -0.10 | 1.00 | 0.01 | 0.02 | 0.59 | 1.00 | -0.01 | 0.02 | -0.48 | 1.00 | -0.01 | 0.02 | -0.48 | 1.00 | -0.00 | 0.02 | -0.21 | 1.00 |
| Dense Urban Poverty | -0.00 | 0.02 | -0.26 | 1.00 | -0.02 | 0.02 | -0.92 | 1.00 | -0.03 | 0.02 | -1.65 | 1.00 | -0.07 | 0.02 | -3.36 | **$1.02 \times 10^{-2}$** | 0.04 | 0.02 | 2.15 | $4.11 \times 10^{-1}$ |
| Extracurriculars | -0.02 | 0.02 | -0.75 | 1.00 | 0.01 | 0.02 | 0.31 | 1.00 | 0.02 | 0.03 | 0.81 | 1.00 | -0.02 | 0.03 | -0.62 | 1.00 | -0.01 | 0.02 | -0.53 | 1.00 |
| Screen Time | -0.04 | 0.03 | -1.63 | 1.00 | -0.04 | 0.03 | -1.37 | 1.00 | -0.11 | 0.03 | -3.62 | **$3.95 \times 10^{-3}$** | -0.06 | 0.03 | -1.92 | $7.09 \times 10^{-1}$ | -0.08 | 0.03 | -2.99 | **$3.64 \times 10^{-2}$** |
| **Replication Sample** | | | | | | | | | | | | | | | | | | | | |
| Intercept | 0.00 | 0.04 | 0.02 | 1.00 | 0.06 | 0.04 | 1.34 | 1.00 | 0.09 | 0.04 | 1.93 | $7.05 \times 10^{-1}$ | 0.09 | 0.04 | 2.09 | $4.82 \times 10^{-1}$ | -0.00 | 0.05 | -0.07 | 1.00 |
| Age | 0.25 | 0.02 | 16.02 | **$3.72 \times 10^{-54}$** | 0.16 | 0.02 | 9.22 | **$7.32 \times 10^{-19}$** | 0.09 | 0.02 | 4.76 | **$2.66 \times 10^{-5}$** | 0.22 | 0.02 | 12.16 | **$4.66 \times 10^{-32}$** | 0.24 | 0.02 | 14.51 | **$7.37 \times 10^{-45}$** |
| Sex | 0.07 | 0.03 | 2.24 | $3.25 \times 10^{-1}$ | 0.01 | 0.04 | 0.29 | 1.00 | -0.10 | 0.04 | -2.59 | $1.26 \times 10^{-1}$ | -0.11 | 0.04 | -2.79 | $6.88 \times 10^{-2}$ | 0.10 | 0.03 | 3.00 | **$3.56 \times 10^{-2}$** |
| ADHD Medications | -0.05 | 0.06 | -0.84 | 1.00 | -0.17 | 0.07 | -2.45 | $1.84 \times 10^{-1}$ | -0.05 | 0.08 | -0.62 | 1.00 | -0.04 | 0.07 | -0.61 | 1.00 | -0.19 | 0.07 | -2.94 | **$4.36 \times 10^{-2}$** |
| Antidepressants | -0.04 | 0.13 | -0.27 | 1.00 | -0.08 | 0.15 | -0.57 | 1.00 | 0.04 | 0.16 | 0.26 | 1.00 | -0.20 | 0.15 | -1.29 | 1.00 | -0.20 | 0.13 | -1.46 | 1.00 |
| Antipsychotics | -0.42 | 0.22 | -1.88 | $7.75 \times 10^{-1}$ | -0.16 | 0.25 | -0.63 | 1.00 | -0.33 | 0.27 | -1.22 | 1.00 | 0.00 | 0.26 | 0.01 | 1.00 | -0.24 | 0.23 | -1.03 | 1.00 |
| Exp-Factor | 0.46 | 0.02 | 24.73 | **$8.94 \times 10^{--121}$** | 0.15 | 0.02 | 7.41 | **$2.24 \times 10^{-12}$** | 0.21 | 0.02 | 9.64 | **$1.53 \times 10^{-20}$** | 0.12 | 0.02 | 5.63 | **$2.59 \times 10^{-7}$** | 0.39 | 0.02 | 19.32 | **$7.49 \times 10^{-77}$** |
| School | 0.02 | 0.02 | 1.01 | 1.00 | 0.02 | 0.02 | 1.08 | 1.00 | 0.01 | 0.02 | 0.30 | 1.00 | 0.05 | 0.02 | 2.29 | $2.86 \times 10^{-1}$ | 0.00 | 0.02 | 0.08 | 1.00 |
| Family Values | -0.11 | 0.02 | -5.72 | **$1.57 \times 10^{-7}$** | -0.05 | 0.02 | -2.27 | $3.06 \times 10^{-1}$ | -0.01 | 0.02 | -0.46 | 1.00 | 0.00 | 0.02 | 0.19 | 1.00 | -0.04 | 0.02 | -2.05 | $5.27 \times 10^{-1}$ |
| Family Turmoil | -0.01 | 0.02 | -0.75 | 1.00 | 0.02 | 0.02 | 0.88 | 1.00 | 0.05 | 0.02 | 2.46 | $1.79 \times 10^{-1}$ | 0.00 | 0.02 | 0.01 | 1.00 | 0.00 | 0.02 | 0.07 | 1.00 |
| Dense Urban Poverty | -0.00 | 0.02 | -0.02 | 1.00 | -0.04 | 0.02 | -1.96 | $6.58 \times 10^{-1}$ | -0.04 | 0.02 | -2.23 | $3.35 \times 10^{-1}$ | -0.01 | 0.02 | -0.52 | 1.00 | 0.01 | 0.02 | 0.70 | 1.00 |
| Extracurriculars | -0.04 | 0.02 | -1.89 | $7.67 \times 10^{-1}$ | -0.03 | 0.02 | -1.39 | 1.00 | -0.04 | 0.03 | -1.68 | 1.00 | -0.03 | 0.03 | -1.20 | 1.00 | -0.04 | 0.02 | -1.83 | $8.77 \times 10^{-1}$ |
| Screen Time | -0.08 | 0.03 | -3.00 | **$3.58 \times 10^{-2}$** | -0.02 | 0.03 | -0.65 | 1.00 | -0.07 | 0.03 | -2.14 | $4.19 \times 10^{-1}$ | -0.06 | 0.03 | -1.78 | $9.74 \times 10^{-1}$ | -0.08 | 0.03 | -2.87 | $5.33 \times 10^{-2}$ |

**Supplementary Table 5. Exp-Factor at baseline assessment is significantly associated with cognition over and above the effects of psychiatric medication use.** Across all five cognitive tasks and across both the discovery and replication sub-samples, the general exposome factor (Exp-Factor) is positively associated with cognition over and above psychiatric medication use (ADHD Medications, Antidepressants and Antipsychotics). Note that random effects for site and family were also included as covariates in these models and that the effects of interest also remained significant in longitudinal models associating the general exposome factor with cognitive performance at the two year follow-up. *Abbreviations:* ADHD: Attention-Deficit/Hyperactivity Disorder.

| *Linear Mixed Effects Models* | $n$ | Picture Vocabulary $\beta_{Exp\text{-}Factor}$ | Picture Vocabulary $p_{bonf}$ | Flanker $\beta_{Exp\text{-}Factor}$ | Flanker $p_{bonf}$ | Picture Sequence Memory $\beta_{Exp\text{-}Factor}$ | Picture Sequence Memory $p_{bonf}$ | Pattern Comparison $\beta_{Exp\text{-}Factor}$ | Pattern Comparison $p_{bonf}$ | Reading Recognition $\beta_{Exp\text{-}Factor}$ | Reading Recognition $p_{bonf}$ |
|---|---|---|---|---|---|---|---|---|---|---|---|
| **Stratification By Biological Sex** | | | | | | | | | | | |
| Female | 5045 | 0.49 | **$8.37 \times 10^{-256}$** | 0.22 | **$2.05 \times 10^{-48}$** | 0.24 | **$2.10 \times 10^{-56}$** | 0.14 | **$1.37 \times 10^{-20}$** | 0.41 | **$7.76 \times 10^{-167}$** |
| Male | 5492 | 0.49 | **$1.47 \times 10^{-272}$** | 0.20 | **$5.68 \times 10^{-38}$** | 0.20 | **$4.39 \times 10^{-43}$** | 0.13 | **$1.05 \times 10^{-15}$** | 0.45 | **$9.60 \times 10^{-196}$** |
| **Stratification By Race** | | | | | | | | | | | |
| White | 6959 | 0.43 | **$5.27 \times 10^{-192}$** | 0.14 | **$1.34 \times 10^{-20}$** | 0.18 | **$3.38 \times 10^{-27}$** | 0.10 | **$1.88 \times 10^{-8}$** | 0.39 | **$1.73 \times 10^{-135}$** |
| Black | 1549 | 0.39 | **$9.39 \times 10^{-49}$** | 0.31 | **$1.92 \times 10^{-15}$** | 0.13 | **$1.65 \times 10^{-5}$** | 0.11 | **$2.70 \times 10^{-3}$** | 0.45 | **$1.25 \times 10^{-46}$** |
| Asian | 236 | 0.41 | **$1.69 \times 10^{-6}$** | 0.12 | 1.00 | 0.17 | $4.39 \times 10^{-1}$ | 0.18 | $3.54 \times 10^{-1}$ | 0.29 | **$2.82 \times 10^{-3}$** |
| AIAN/NHPI | 66 | 0.69 | **$8.89 \times 10^{-4}$** | 0.54 | **$2.25 \times 10^{-2}$** | 0.34 | $2.29 \times 10^{-1}$ | 0.07 | 1.00 | 0.38 | $4.34 \times 10^{-1}$ |
| Other/Mixed | 1727 | 0.44 | **$2.09 \times 10^{-80}$** | 0.19 | **$3.17 \times 10^{-13}$** | 0.17 | **$2.94 \times 10^{-13}$** | 0.13 | **$4.88 \times 10^{-7}$** | 0.42 | **$3.03 \times 10^{-59}$** |

**Supplementary Table 6.** Stratified analyses by biological sex and racial identification. Results are shown for the Exp-Factor coefficient ($\beta_{\text{Exp-Factor}}$) from models associating the exposome with cognitive functioning across five tasks while accounting for covariates (age, family, site, and the specific exposome sub-factors). Biological sex was also included as a covariate for the race stratification analysis in line with our primary models. We observed consistent positive associations between Exp-Factor and cognitive functioning for nearly all cognitive tasks in all groups, with only a small subset of associations not passing Bonferroni correction for multiple comparisons ($p_{\text{bonf}}$). To ensure that there was a sufficient sample size available for all racial categories, racial identifications of “Other” and “Mixed” were combined and all stratified analyses were performed on the whole sample at baseline assessment.

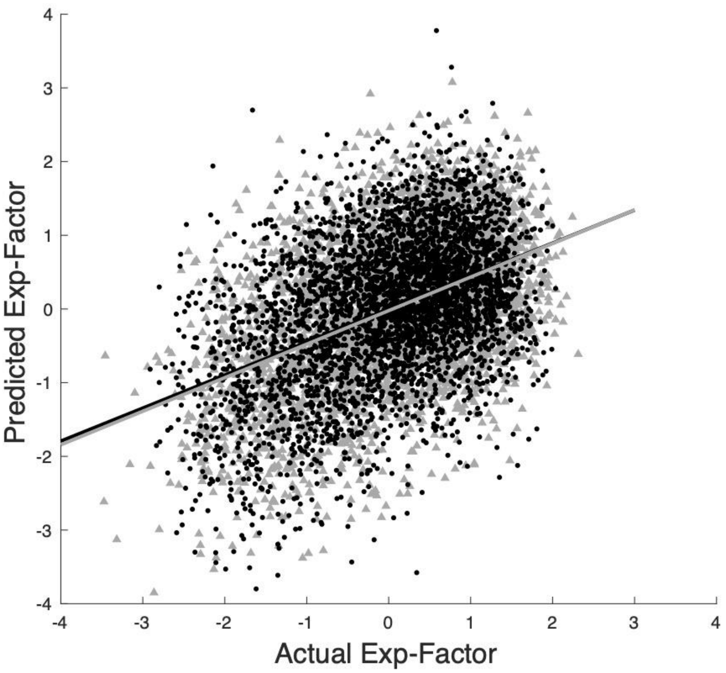


**Supplementary Figure 3.** Associations between true exp-factor and model-generated exp-factor using exposome scores generated independently in the discovery and replication samples rather than from the full sample of participants. Confirming that our results were not driven by leakage across samples, we found nearly identical results (discovery: $r = 0.440$, $p<0.001$, 95% CI: [0.41, 0.47]; replication: $r = 0.460$, $p<0.001$, 95% CI: [0.43, 0.49]).

| | Discovery | | | | Replication | | | |
|---|---|---|---|---|---|---|---|---|
| *Prediction Accuracy* | *r* | *p* | AIC | BIC | *r* | *p* | AIC | BIC |
| **Picture Vocabulary** | | | | | | | | |
| Exp-Factor | 0.18 | $5.42 \times 10^{-17}$ | $1.8908 \times 10^{4}$ | -25.6472 | 0.15 | $9.75 \times 10^{-12}$ | $1.8706 \times 10^{4}$ | -25.6492 |
| PFN Topography | 0.11 | $3.88 \times 10^{-7}$ | $2.0389 \times 10^{6}$ | $7.7267 \times 10^{6}$ | 0.09 | $1.13 \times 10^{-4}$ | $2.0387 \times 10^{6}$ | $7.7151 \times 10^{6}$ |
| Exp-Factor + PFN Topography | 0.12 | $6.28 \times 10^{-8}$ | $2.0389 \times 10^{6}$ | $7.7267 \times 10^{6}$ | 0.09 | $3.34 \times 10^{-5}$ | $2.0387 \times 10^{6}$ | $7.7151 \times 10^{6}$ |
| **Flanker** | | | | | | | | |
| Exp-Factor | 0.11 | $2.27 \times 10^{-7}$ | $1.9389 \times 10^{4}$ | -26.1050 | 0.10 | $2.33 \times 10^{-6}$ | $1.9171 \times 10^{4}$ | -26.0975 |
| PFN Topography | 0.08 | $1.97 \times 10^{-4}$ | $2.0394 \times 10^{6}$ | $7.7267 \times 10^{6}$ | 0.08 | $1.60 \times 10^{-4}$ | $2.0392 \times 10^{6}$ | $7.7151 \times 10^{6}$ |
| Exp-Factor + PFN Topography | 0.08 | $1.42 \times 10^{-4}$ | $2.0394 \times 10^{6}$ | $7.7267 \times 10^{6}$ | 0.08 | $1.25 \times 10^{-4}$ | $2.0392 \times 10^{6}$ | $7.7151 \times 10^{6}$ |
| **Picture Sequence Memory** | | | | | | | | |
| Exp-Factor | 0.16 | $2.12 \times 10^{-13}$ | $1.9780 \times 10^{4}$ | -26.4773 | 0.18 | $3.94 \times 10^{-17}$ | $1.9562 \times 10^{4}$ | -26.4738 |
| PFN Topography | 0.11 | $1.25 \times 10^{-6}$ | $2.0398 \times 10^{6}$ | $7.7267 \times 10^{6}$ | 0.13 | $5.08 \times 10^{-9}$ | $2.0396 \times 10^{6}$ | $7.7151 \times 10^{6}$ |
| Exp-Factor + PFN Topography | 0.11 | $3.84 \times 10^{-7}$ | $2.0398 \times 10^{6}$ | $7.7267 \times 10^{6}$ | 0.13 | $8.91 \times 10^{-10}$ | $2.0396 \times 10^{6}$ | $7.7151 \times 10^{6}$ |
| **Pattern Comparison** | | | | | | | | |
| Exp-Factor | 0.05 | 0.014 | $1.9562 \times 10^{4}$ | -26.2696 | 0.13 | $3.41 \times 10^{-9}$ | $1.9357 \times 10^{4}$ | -26.2763 |
| PFN Topography | 0.06 | 0.009 | $2.0396 \times 10^{6}$ | $7.7267 \times 10^{6}$ | 0.04 | 0.047 | $2.0394 \times 10^{6}$ | $7.7151 \times 10^{6}$ |
| Exp-Factor + PFN Topography | 0.06 | 0.008 | $2.0396 \times 10^{6}$ | $7.7267 \times 10^{6}$ | 0.05 | 0.036 | $2.0394 \times 10^{6}$ | $7.7151 \times 10^{6}$ |
| **Reading Recognition** | | | | | | | | |
| Exp-Factor | 0.06 | 0.003 | $1.9162 \times 10^{4}$ | -25.8895 | 0.12 | $3.52 \times 10^{-8}$ | $1.8942 \times 10^{4}$ | -25.8765 |
| PFN Topography | 0.04 | 0.074 | $2.0392 \times 10^{6}$ | $7.7267 \times 10^{6}$ | 0.05 | 0.018 | $2.0392 \times 10^{6}$ | $7.7151 \times 10^{6}$ |
| Exp-Factor + PFN Topography | 0.04 | 0.062 | $2.0389 \times 10^{6}$ | $7.7267 \times 10^{6}$ | 0.05 | 0.013 | $2.0390 \times 10^{6}$ | $7.7151 \times 10^{6}$ |

**Supplementary Table 7. Comparison of longitudinal models relating exposome and PFN topography to cognitive functioning across domains.** To compare multivariate associations among exposome scores, personalized functional brain network topography and cognitive functioning, we trained linear ridge regression models to predict cognitive performance assessed two years later when children were 11-12 years old. The first model type (“Exp-Factor”) used only a participant’s general exposome score, while the second model type (“PFN Topography”), reported in our prior work (Keller et al., 2022), used each participant’s multivariate pattern of personalized functional network topography. The third model type (“Exp-Factor + PFN Topography”) used a combination of the features in the first and second model types. Correlations between true cognitive performance and model-generated cognitive performance (*r*) were significant for all model types and highest for the predictions of General Cognition. Model comparison using the Akaike Information Criterion (AIC) and Bayesian Information Criterion (BIC), indices that account for differences in the number of features across models, reveals that the Exp-Factor model is the most parsimonious.

| Scale | Sub-Score | Items |
| --- | --- | --- |
| School Risk and Protective Factors Survey | school_Enjoy | school_12_y, school_15_y (REVERSED), school_3_y |
| | school_FeelInvolved | school_10_y, school_5_y, school_2_y, school_6_y |
| | school_GoodGrades | school_8_y, school_17_y (REVERSED), school_9_y |
| | school_PositiveFeedback | school_7_y, school_4_y |
| youth-report Family Environment Scale | family_probs_youth_BadKey | fes_youth_q1, fes_youth_q3, fes_youth_q5, fes_youth_q6, fes_youth_q8 |
| | family_probs_youth_GoodKey | fes_youth_q2, fes_youth_q4, fes_youth_q7, fes_youth_q9 |
| parent-report Family Environment Scale | family_probs_parent_BadKey | fam_enviro1_p, fam_enviro3_p, fam_enviro5_p, fam_enviro6_p, fam_enviro8_p |
| | family_probs_parent_GoodKey | fam_enviro2r_p, fam_enviro4r_p, fam_enviro7r_p, fam_enviro9r_p |
| Parent Monitoring Survey | parent_monitor | parent_monitor_q1_y, parent_monitor_q2_y, parent_monitor_q3_y, parent_monitor_q4_y, parent_monitor_q5_y |
| Neighborhood Safety/Crime Survey | nbrhd_safety_parent | neighborhood1r_p, neighborhood2r_p, neighborhood3r_p |
| Community Risk and Protective Factors | substance_risk | su_risk_p_1, su_risk_p_2, su_risk_p_3, su_risk_p_4, su_risk_p_5 |
| Mexican-American Cultural Values Scale | mexAm_Religiosity | mex_american25_p, mex_american15_p, mex_american11_p, mex_american20_p, mex_american6_p, mex_american1_p, mex_american28_p |
| | mexAm_CaringSecurity | mex_american16_p, mex_american13_p, mex_american7_p, mex_american21_p, mex_american8_p, mex_american26_p, mex_american9_p, mex_american23_p, mex_american14_p, mex_american17_p, mex_american12_p |
| | mexAm_FamilyImage | mex_american18_p, mex_american4_p, mex_american27_p, mex_american3_p, mex_american2_p, mex_american22_p |

| | | |
|---|---|---|
| | mexAm_Independence | mex_american10_p, mex_american5_p, mex_american24_p, mex_american19_p |
| Parental Rules on Substance Use | parent_rules | parent_rules_q1, parent_rules_q4, parent_rules_q7 |
| Sports and Activities Involvement Questionnaire | activities_Feminine | sai_p_activities___24, sai_p_activities___0, sai_p_activities___26, sai_p_activities___22, sai_p_activities___25, sai_p_activities___9, sai_p_activities___6, sai_p_activities___23, sai_p_activities___8, sai_p_activities___17, sai_p_read, sai_p_activities___27 |
| | activities_Masculine | sai_p_activities___5, sai_p_activities___2, sai_p_activities___1, sai_p_activities___13, sai_p_activities___20, sai_p_activities___12, sai_p_activities___14 |
| | activities_Other | sai_p_activities___18, sai_p_activities___28, sai_p_lmusic, sai_p_activities___21, sai_p_activities___19, sai_p_activities___3, sai_p_activities___11, sai_p_activities___16, sai_p_activities___15, sai_p_activities___7, sai_p_activities___4, sai_p_activities___10 |
| Traumatic Brain Injury sum scores | TBIs | tbi_1, tbi_2, tbi_3, tbi_4, tbi_5, tbi_6o, tbi_7a |
| Youth Substance Use Attitudes Questionnaire | peer_deviance | peer_deviance_1, peer_deviance_2, peer_deviance_3, peer_deviance_4, peer_deviance_8, peer_deviance_9 |
| hours of screen time on various devices | screen_time | screentime_11_wknd_br, screentime_5_wkdy_br, screentime_6_wkdy_br, screentime_12_wknd_br, screentime_14_wknd_br, screentime_8_wkdy_br, screentime_2_wkdy_br, screentime_8_wknd_br, screentime_1_wkdy_br, screentime_7_wknd_br |

| | | |
|---|---|---|
| address-based geographic data | poverty | reshist_addr1_coi_se_public, reshist_addr1_coi_se_single, reshist_addr1_adi_sp, reshist_addr1_adi_perc, reshist_addr1_coi_c5_coi_nat (REVERSED), reshist_addr1_coi_se_povrate, reshist_addr1_adi_unemp, reshist_addr1_svi_th2_20142018, reshist_addr1_adi_in_dis, reshist_addr1_coi_r_ed_nat (REVERSED), reshist_addr1_adi_ncar, reshist_addr1_coi_se_emprat (REVERSED), reshist_addr1_coi_se_mhe (REVERSED), reshist_addr1_svi_sin_20142018, reshist_addr1_coi_se_occ (REVERSED), reshist_addr1_svi_dis_20142018, reshist_addr1_svi_emp_20142018, reshist_addr1_coi_he_vacancy, reshist_addr1_coi_he_food, reshist_addr1_coi_c5_he_met (REVERSED), reshist_addr1_adi_mortg (REVERSED), reshist_addr1_coi_r_he_nat (REVERSED), reshist_addr1_svi_veh_20142018, reshist_addr1_coi_ed_schpov, reshist_addr1_adi_rent (REVERSED), reshist_addr1_adi_edu_h (REVERSED), reshist_addr1_opat_kfrpp_p100 (REVERSED), reshist_addr1_svi_min_20142018, reshist_addr1_coi_se_home (REVERSED) |
| | dense_suburban | reshist_addr1_no2_2016_aavg, reshist_addr1_no2_2016_max, reshist_addr1_coi_ed_prxece, reshist_addr1_coi_he_walk, reshist_addr1_urban_area (REVERSED), reshist_addr1_coi_he_green, reshist_addr1_svi_mob_20142018 (REVERSED), reshist_addr1_coi_ed_prxhqece, reshist_addr1_popdensity, reshist_addr1_d1a, reshist_addr1_adi_work_c |

| | | |
|---|---|---|
| | crowding_and_crime | reshist_addr1_adi_crowd, reshist_addr1_p1tot, reshist_addr1_svi_crwd20142018, reshist_state_immigrant_factor (REVERSED), reshist_addr1_pm25_2016_min, reshist_addr1_coi_he_hlthins (REVERSED) |
| | dry_heat | reshist_addr1_scanweekvpd_t3, reshist_addr1_scanweekvpd_t2, reshist_addr1_scanweekvpd_t4, reshist_addr1_scanweekvpd_t1, reshist_addr1_scanweekvpd_t5, reshist_addr1_scanweekvpd_t0, reshist_addr1_scanweekvpd_t6, reshist_addr1_scanweektemp_t0 |
| | traditional_south_and_midwest | reshist_state_racism_factor, reshist_state_sexism_factor, reshist_state_mj_laws, reshist_addr1_coi_he_heat, reshist_addr1_pm252016aa |
| | avg_air_pollution | reshist_addr1_coi_he_pm25, reshist_addr1_no2, reshist_addr1_pm25_2016daysepa (REVERSED), reshist_addr1_pm25, reshist_addr1_pm25_2016_max (REVERSED) |
| | ozone | reshist_addr1_o3_2016_annavg, reshist_addr1_coi_he_ozone (REVERSED), reshist_addr1_coi_ed_college, reshist_addr1_o3_2016_min, reshist_addr1_no2_2016_min, reshist_addr1_coi_he_rsei (REVERSED), reshist_addr1_elevation (REVERSED) |
| | retirement_and_group_living | reshist_addr1_svi_hous20142018, reshist_addr1_svi_th4_20142018, reshist_addr1_svi_grp_20142018, reshist_addr1_svi_17_20142018 (REVERSED) |

**Supplementary Table 8.** Items included in each subscore derived for the longitudinal bifactor analysis.

| Factor | Explained Common Variance (SS) | Omega | Omega-hierarchical (OmegaH) | H | Factor Determinacy |
|---|---|---|---|---|---|
| Exp-Factor | 0.364 | 0.868 | 0.457 | 0.858 | 0.894 |
| School | 0.189 | 0.865 | 0.111 | 0.745 | 0.859 |
| Family Values | 0.144 | 0.868 | 0.068 | 0.739 | 0.874 |
| Family Turmoil | 0.078 | 0.868 | 0.049 | 0.504 | 0.704 |
| Dense Urban Poverty | 0.157 | 0.868 | 0.146 | 0.665 | 0.777 |
| Extracurriculars | 0.052 | 0.868 | 0.033 | 0.381 | 0.619 |
| Screen Time | 0.020 | 0.868 | 0.011 | 0.190 | 0.431 |

**Supplementary Table 9. Bifactor indices for exposome model.** *Abbreviations*: Explained Common Variance (SS) = Explained common variance of a specific factor with respect to itself; H = replicability for all factors given standardized factor loadings; OmegaH =  hierarchical omega reliability estimate.

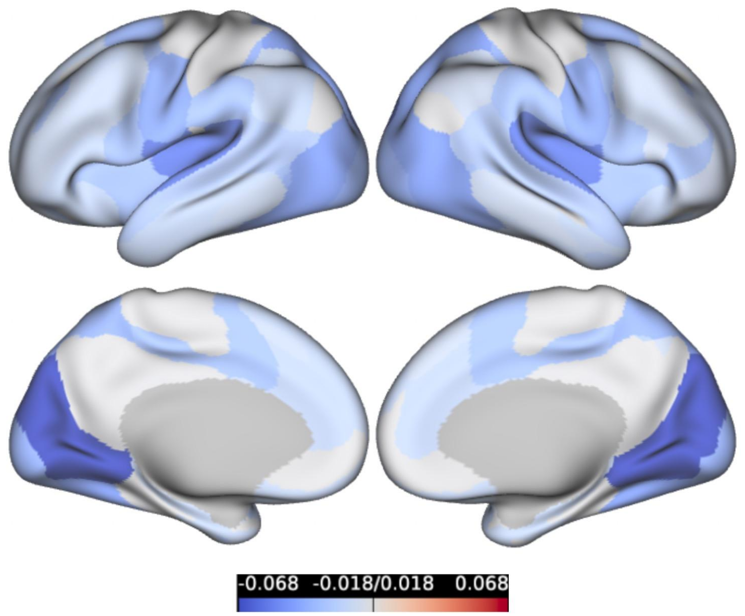


**Supplementary Figure 4. Differences in prediction accuracy by PFN between exposome and cognition models.** To identify differences in the spatial distribution of prediction accuracy across PFNs between models, we subtracted the map of prediction accuracy by PFN for a model predicting general exposome factor from the map of prediction accuracy by PFN for a model predicting general cognition. Prediction accuracy is defined as the correlation between the observed general exposome factor (or observed general cognition) and the estimated general exposome factor (or estimated general cognition) from ridge regression models trained on PFN functional topography. The very small difference scores between these two maps suggest that the spatial patterning of prediction accuracies by PFN do not vary substantially between these two models, and indeed the correlation between the two maps was high ($r$ = 0.933, $p = 4.842 \times 10^{-8}$). The negative values for these difference scores suggest that for all PFNs, models predicting general cognition outperform models predicting general exposome scores.
